# Lineage domains and cytoskeletal cables organize a cellular square grid in a crustacean

**DOI:** 10.1101/2025.08.31.673345

**Authors:** Beatrice L. Steinert, Leo Blondel, Chandrashekar Kuyyamudi, Evangelia Stamataki, Anastasios Pavlopoulos, Cassandra G. Extavour

## Abstract

To build tissues, and ultimately functional bodies, cells in early embryos must arrange into specific patterns. In most animals, epithelial tissues exhibit a predominantly hexagonal space-packing geometry. However, in many species of the largest group of crustaceans, the Malacostraca, the embryonic epithelium takes on the striking form of a grid made up of predominantly square cells, sequential rows of which establish the adult segmented body plan. After square cells emerge, their organization appears to be maintained by specific cell division patterns. However, the mechanisms that initially generate square cells from hexagonal precursors are unknown. Here we address this problem by combining long-term multiview lightsheet microscopy, immunohistochemistry, laser ablation, and pharmacological perturbation. We show that in the emerging model crustacean *Parhyale hawaiensis* this highly unusual grid geometry is first initiated from two perpendicular axes that are established sequentially according to different cellular mechanisms. The first axis arises dorso-ventrally at a tensile lineage compartment boundary, while the second emerges at the anterior-posterior axis along the ventral midline through lineage-independent cell intercalation driven by tensile actomyosin cables. We show that these midline cables are necessary for organizing square-cell packing as well as for proper expression of the segmentation gene *engrailed*. Our findings show that both cell lineage-specific behaviors, as well as lineage-independent supracellular structures, are required to establish square grid epithelial organization and a segmented body plan.

## Introduction

The organization of cells and tissues during development is fundamental for multicellular organisms. During embryogenesis, cells must be shaped at both the individual and collective tissue levels to form functional structures at every stage of ontogeny. It has been appreciated since the late nineteenth century that mechanical forces and geometric properties of cells and tissues play a central role in morphogenesis, the generation of form during development^1^. Despite the predominant focus on the role of genes in tissue patterning during the last half century^2–5^, recent decades have witnessed a resurgence of interest in the mechanics of development at the cell and tissue scales of biological organization^6–8^. Harnessing techniques for live imaging and computational image processing and annotation, recent work has begun to couple mechanical and genetic contributions to tissue organization during embryogenesis^9^. However, these investigations have been largely limited to a small number of model laboratory organisms, giving us a narrow phylogenetic sampling.

The arthropods, which include insects and crustaceans, are and always have been the most speciose group of animals on earth^10^. All arthropods have segmented body plans that are established early in embryonic development^11^. Segmentation is evident in embryonic ectodermal epithelia^12^, tissues in which cells are tightly adhered to one another in a monolayer sheet and apical cell faces form a confluent network of polygons^13^. Most epithelia that have been studied to date are those in which constituent cells exhibit predominantly hexagonal packing with three-way vertices, which is the geometry with minimal surface energy and maximal space-filling efficiency^14–18^. During segmentation in the fruit fly *Drosophila melanogaster*^19,20^ and the milkweed bug *Oncopeltus fasciatus*^21^, a small number of cells at segment boundaries briefly stray from this geometry to give rise to an increased number of 4-way vertices, suggesting an altered mechanical state. However, most epithelial cells retain their hexagonal shapes throughout segmentation and early morphogenesis^22–24^. In *D. melanogaster*, maintaining boundaries between embryonic segments requires tensile cables of actin and non-muscle myosin (myosin 2), which are mechanically coupled across the tissue and prevent cell mixing across the boundary^19,24^.

In contrast, in the early embryonic ectodermal epithelium of malacostracan crustaceans (including shrimp, lobsters, isopods and amphipods) cells are packed into a grid of square cells^25,26^ arranged into highly ordered rows and columns^27,28^. Expression of the segment polarity gene *engrailed* is initiated precisely every four rows of the square grid^29^. This emergence and maintenance of square cells across the tissue is particularly striking given that higher surface energy is predicted to be required for this space-packing geometry, and that 4-way vertices, which are essential for square-packing, are unstable in vertex models of epithelia^30^. The amphipod crustacean *Parhyale hawaiensis* is a powerful model for deepening our understanding of how local lineage relationships and physical parameters of individual cells can result in correct tissue morphogenesis and body patterning^31^. In contrast with *D. melanogaster*, in which early patterning is set in motion within a syncytial embryo^32^, the early *P. hawaiensis* embryo cleaves holoblastically in a highly stereotyped manner that defines the cellular progenitors of all three germ layers and the germ line as individual blastomeres of the 8-cell embryo^27^. From this stage onwards, the three blastomeres that will give rise to the ectoderm–the descendants of which make up the square-packing epithelium–can be tracked and their clones spatially mapped^27,33^.

Here we uncover the cellular mechanisms underlying morphogenesis of this highly ordered tissue in *P. hawaiensis*. Previous work in this amphipod suggests that after square cells appear, their packing into a grid is maintained by cell divisions that occur in bilaterial, palindromic medial-to-lateral mitotic waves during the phase when grid rows duplicate along the anterior-posterior axis to establish clonally related parasegments^34^. However, the mechanisms that ensure that 4-way vertices, square cells, and the tissue-wide arrangement of cells into rows and columns emerge in the first place, remain unknown.

To address this problem, we used long-term, multiview lightsheet and confocal microscopy, and imaged *P. hawaiensis* embryos from the 16-cell stage through to the first emergence of square cells and the square grid (Fig. 1a). We used semi-automated cell tracking to trace the ectodermal lineages and map their clonal domains onto the quantitative dynamics of cell shape within the expanding epithelium. We show that square-packing geometry emerges along two axes, corresponding with the anterior-posterior and dorsal-ventral body axes, through both lineage-dependent and lineage-independent morphogenetic mechanisms within the tissue. We further show that supracellular actomyosin cables form at these axes, and that the cables are under locally-generated tension higher than that of the surrounding tissue. Finally, we provide evidence that these cables function as a mechanical organizer of this tissue.

**Figure 1.**
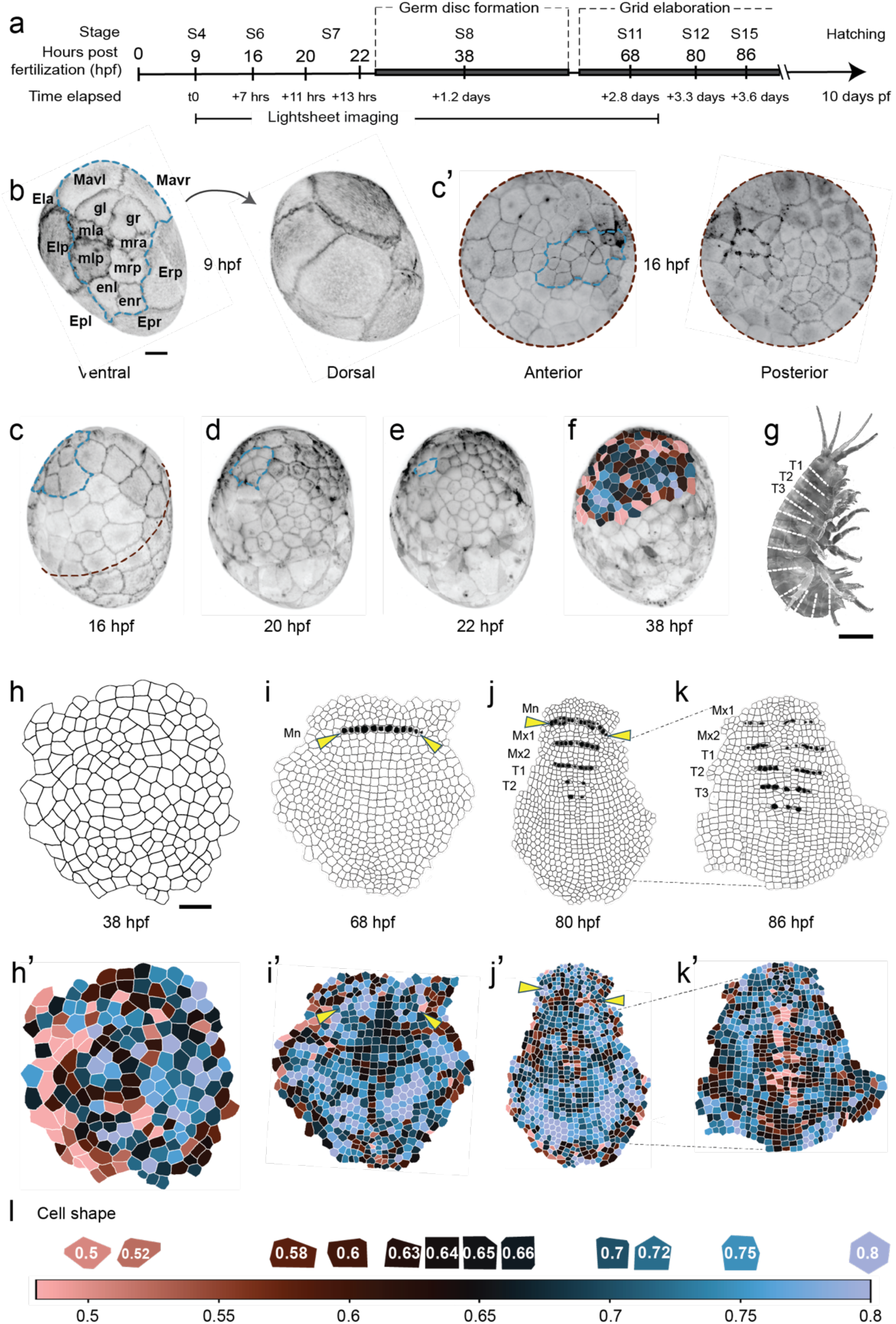
Formation of a cellular square grid in the *Parhyale hawaiensis* embryonic ectoderm. **a** Developmental timeline from fertilization to the hatching of juveniles with the window of lightsheet microscope imaging indicated at the bottom. Stages according to Browne and colleagues^29^. **b-f** Stills selected from a time-lapse of the first 1.5 days of development in a *P. hawaiensis* embryo that was live-imaged with a multiview lightsheet microscope; cell membranes marked with *Lyn-EGFP*. Micrographs are maximum intensity projections of a volume constructed by fusing 3D image stacks from all imaging angles, and are displayed either from a fixed perspective (b-f) or as conformal surface maps (c’). **b** Ventral (left) and dorsal (right) views of the 16-cell stage. The first cell divisions in *P. hawaiensis* are highly stereotypic and segregate the germ layers and germ line (gl and gr) by this stage. The eight micromeres and two macromeres enclosed within the blue dotted line are the mesoderm (ml, mr, Mav), endoderm (en), and germ-line (g) precursors, the descendants of which delaminate from the surface during gastrulation (c-e) leaving at the surface a continuous sheet of epithelial ectoderm (f). The remaining six macromeres (Epr, Epl, Era, Erp, Ela, Elp) will give rise to all ectodermal cells. **c-e** From approximately 16-22 hours post fertilization (hpf), cells begin reducing in size and migrate to the anterior pole, condensing into a germ disc (embryonic rudiment) sitting atop larger yolk-filled cells. **c’** Anterior (left) and posterior (right) conformal maps of the same image volume displayed in c. These 2D surface projections allow for simultaneous visualization of cell shapes and relationships that make up the embryo’s curved surface. The maroon dotted line in c corresponds to those along map edges in c’. **f** By approximately 38 hpf, the surface of the germ disc is an epithelium made up entirely of ectoderm (colored cells). **g** An adult female *P. hawaiensis.* White lines indicate parasegmental boundaries with corresponding morphological segment identities. **h-k** Time series of fixed embryos in which the ectoderm was dissected away from the yolk and physically flattened to make surface cell geometries visible. A grid of four-sided cells emerges within the expanding germ disc. The transition from hexagonal to square cells (**h’-k’**) progresses from anterior to posterior and laterally left and right (**i**). A row of cells expressing the segmentation gene *engrailed* appears in the prospective mandibular segment (Mn) around 68 hpf (arrowheads). **j, k** From 80 hpf onwards, progressing from anterior to posterior, nuclear-localized Engrailed protein is detectable in cells of the anterior-most cell row in each parasegment as it is generated via row-duplicating cell divisions from the midline to lateral regions. Dotted lines in j indicate the region of the embryo displayed in k. **l** Cell shape descriptor used to analyze square grid morphogenesis in this study. While four-sided cells can be either square or rectangular, for simplicity we describe them as square throughout the text, and our cell shape descriptor gives square cells a metric value of 0.64. Mx1: first maxillary segment; Mx2: second maxillary segment; T1: first trunk segment; T2: second trunk segment; T3: third trunk segment.

## Results

### Ectodermal grid formation

The first seven cleavages of the early *P. hawaiensis* embryo are holoblastic, and the progenitors of all three germ layers and the germ line are established at the 8-cell stage^27,29,33,35^. Beginning at approximately 20 hours post fertilization (“hpf” hereafter), cells that will contribute to the embryo migrate towards the anterior pole and coalesce into an embryonic rudiment referred to as a “germ disc,” positioned atop yolk and surrounded by extraembryonic tissue (Fig. 1d). By 38 hpf, after gastrulation (Fig. 1c-e, blue dotted line^33,36^), the ventral surface of this disc (Fig. 1f, h) consists of a monolayer of ectodermal cells^15^. By 68 hpf, the cells in the germ disc have begun aligning into rows and columns and transforming their apical faces into squares (Fig. 1i, j). This square grid sets up the logic of segmentation of the adult body plan (Fig. 1g), with the conserved segment polarity gene *engrailed* becoming expressed along its rows (Fig. 1i-k)^29,37^.

To visualize this epithelial morphogenesis and its relationship to the ectodermal lineages established at the 16-cell stage (Fig. 1b outside of blue dotted line, Supplementary Fig. 1a), we generated volumetric, long-term timelapse images of *P. hawaiensis* embryos injected with nuclear marker *H2B-mCherry* and plasma membrane marker *Lyn-EGFP* mRNA reporter constructs using multiview lightsheet microscopy (Fig. 1a, Movies 1-2). We used a combination of manual and semi-automated methods to annotate cell lineage, cell positions and mitotic activity throughout the entire embryo in frames captured every ten minutes over more than 60 hours of development. We captured the period starting from the eight cell stage at 9 hpf (Stage 4 as per Browne and colleagues^29^) through to Stage 11^29^ at approximately 70-74 hpf (Fig. 1a). By the end of the capture period, the ectodermal grid of square cells that is the subject of this study is clearly established (Fig. 1i, i’). We performed this annotation for three embryos, all of which hatched successfully following imaging (Supplementary Table 1).

We wished to determine whether and how cell lineage, revealed by the tracking described above, was related to the cell shape dynamics and rearrangements that establish the square grid (Movie 3). However, in the later time-points of these datasets, when the germ disc has expanded over the gradual curve of the embryo’s surface, maximum intensity projections of the image volume obscure apical cell morphologies and challenge accurate quantification of cell shape dynamics. To solve this problem, we conformally mapped the 3D surface of the embryo onto a 2D plane using the tissue cartography tool ImSAnE^34,38^ (Fig. 1c’). To visualize the grid at even later stages when the square-cell packing geometry has extended throughout the expanding ectoderm, we dissected fixed embryos from the underlying yolk and physically flattened the ectoderm into the 2D plane (Fig. 1h-k). The surface curvature of the unperturbed tissue at these stages is gradual enough that, unlike in some large vertebrate embryos^39^, flattening does not require incisions and does not deform cell shapes. Using images of these flattened epithelia, we quantified the dynamics of cell shape change in the ectoderm, as well as the relationship of these dynamics to the lineage relationships of the constituent ectodermal cells, from the 16-cell stage through to the establishment of the grid of square cells that serves as the template for the segmented body plan.

### Square-packing geometry initiates at a lineage compartment boundary

Previous work has suggested that an ectodermal lineage compartment boundary runs perpendicular to the anterior-posterior axis of the *P. hawaiensis* embryo^40^. Given the early stereotypic cleavage patterns of this embryo, we asked whether lineage could contribute to the initiation of square-grid morphogenesis. We tracked all six of the 16-cell ectodermal lineages (Era, Ela, Erp, Elp, Epr, Epl; Supplementary Fig. 1a; nomenclature as per Gerberding and colleagues^27^) in the lightsheet animations, enabling us to delineate the dynamic lineage territories within the developing germ disc. We observed that a strict boundary emerged immediately after gastrulation between the anterior (Eva, Ela, grey in Fig. 2a’) and posterior (Erp, Elp, Epr, Epl, striped in Fig. 2a’) lineages, and that clones from each domain never crossed this boundary (Supplementary Fig. 1b, c). This boundary, which we call the initiation line (IL, arrowheads in Fig. 2a), straightened into a line perpendicular to the anterior-posterior axis by 68 hpf (Fig. 2b). The IL was made up of over 30% 4-way vertices from 38-56±2 hpf, and of over 40% 4-way vertices by 62±2 hpf (Fig. 2c). This shows that the first emergence of large numbers of 4-way vertices, required for square cell formation, is coincident with the formation of the IL, which is an ectodermal compartment boundary.

**Figure 2.**
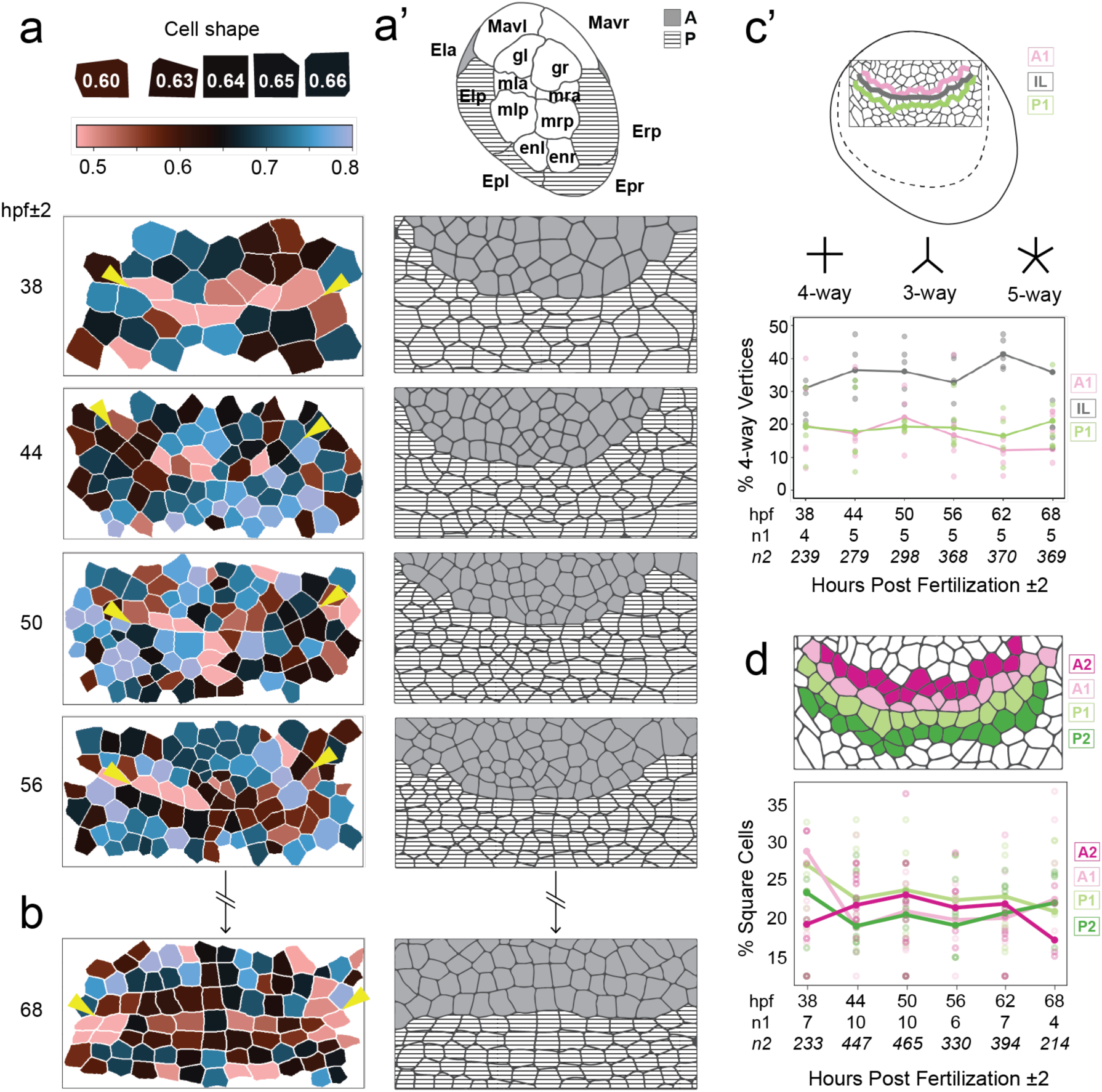
Square cell geometry initiates along the dorso-ventral axis at the initiation line, which is an ectodermal lineage compartment boundary. **a-b** 38-68±2 hpf manually flattened germ discs segmented to identify cell membranes, with cells colored according to their shape descriptor value. What we call the “initiation line” (IL; arrowheads in the left column), corresponds to the boundary of the anterior (A: grey) and posterior (P: striped) lineage compartments shown in a’. **a’** Stills from a lightsheet time-lapse that were conformally mapped onto 2D, showing descendants of ectodermal precursor macromeres (grey: anterior ectoderm founders Era, Ela; striped: posterior ectoderm founders Erp, Elp, Epr, Epl) The boundary between anterior and posterior clones begins as a jagged curve at 38±2 hpf and straightens out by 68±2 hpf. **c** Top: Schematic shows the analyzed region (box) within the germ disc (dotted line). Bottom: Percentage of 4-way vertices at the IL (gray), the equivalent line one cell anterior (A1, light pink), and one cell posterior (P1, light green). **d** Top: Cell rows analyzed in four different locations relative to the IL: immediately adjacent to the IL to the anterior (A1: light pink), immediately adjacent to the IL to the posterior (P1: light green), one cell diameter anterior to the IL (A2: dark pink) and one cell diameter posterior to the IL (P2: dark green). Bottom: plot showing percentage of square cells (shape descriptor value ⩾ 0.60 or ⩽0.66). Anterior is up in all images. Sample sizes in c and d indicated by n1: number of embryos and n2: number of vertices.

### 4-way cellular vertices first emerge in the local vicinity of the IL

Given that 4-way vertices are essential for square cell-packing, and that such vertices first appear in large numbers at the IL, we next characterized the emergence of square cells at the IL. From 38-68 hpf, the cells near the IL exhibited extensive changes in shape and arrangement (Fig. 2a). The occurrence of cell rearrangement at the IL was evident by the dynamics of the four 16-cell stage posterior ectodermal lineage domains (Erp, Elp, Epl, Epr; Supplemental Fig. 1a). Whereas in the 26±2 hpf germ disc the Elp and Erp domains were on opposite sides (left and right) of the germ disc, separated from each other primarily by descendants of the anterior Era and Ela clones (Supplementary Fig. 1b, c, top row), they ultimately became separated by a single column of posterior cells descended from one or both of Epl and Epr (Supplementary Fig. 1b, c, bottom row). We also observed cell proliferation throughout the germ disc, including directly at the IL and in the anterior and posterior compartments, throughout this time window (Supplementary Fig. 2, Supplementary Table 2).

To quantitatively describe the cell shape dynamics, including the transient changes in edge numbers that result from a neighboring cell division and the instability of 4-way vertices^30^, we developed a cell shape descriptor metric, defined as the ratio of the area of a cell to the area of the bounding circle of that cell. For this metric, values range from 0.5 to 1, where higher values are closer to hexagons (Fig. 1l). We defined “square cells” as those with values in the range of 0.6-0.66 (darkest color in Fig. 1l). We analyzed the changes in shape in cells along the IL (Fig. 2d, A1 and P1), and in cells one cell diameter away from the IL in both the anterior and posterior compartments (Fig. 2d, A2 and P2). At 38 hpf, the rows immediately anterior (A1) and posterior (P1) to the IL had more square cells (> 25%) than other measured regions (Fig. 2d). After 38 hpf, P1 and the row one cell diameter anterior to the IL (A2) had the highest proportion of square cells (20-25%) until 62 hpf, but by 68 hpf the proportion of square cells in A2 dropped to less than 20% (Fig. 2d). The proportion of square cells immediately anterior to the IL (A1), and one cell diameter away posterior to the IL (P2), was similar in both regions from 44 to 68 hpf (Fig. 2d). The proportion of square cells in both regions increased by 68 hpf, comprising the highest proportion of square cells of all measured regions. This gradual and local increase in square cells near the IL, suggests that this lineage boundary initiates and maintains the transition to square-packing geometry along the dorsal-ventral axis.

### A simulation of cell sorting can explain the emergence of the IL as a straight boundary

Given that the IL is a lineage compartment boundary, we asked whether and how lineage might contribute to the emergence of this first grid axis, which becomes increasingly straight from 26-68 hpf (Fig. 2a, Supplementary Fig. 1b, c). To test the hypothesis that minimization of interfacial free energy at the boundary between cells of distinct ectodermal lineages might be responsible for boundary formation, we used CompuCell3D^41–43^ to simulate cell sorting within the *P. hawaiensis* ectoderm. To obtain biologically realistic parameters for this simulation, we first segmented lightsheet images at 26 hpf, the first time point after the completion of gastrulation, when the surface epithelium is made up entirely of ectoderm cells (Figs. 2a, 3d, Supplementary Fig. 1b). We then quantified the number of cell divisions within each of the six 16-cell stage ectodermal lineages (Fig. 3a) up to 20 hours after this time point (46 hpf). The membrane segmentation and lineage proliferation rates, as well as increased contact energy between the anterior and posterior lineages (Fig. 3b’), were the initial conditions for the simulation. When run, the model consistently (n = 10 runs) produced a straight anterior-posterior compartment boundary as well as a clonal domain mapping like those observed in the actual tissue (Fig. 3e; compare with Supplementary Fig. 1b, c, bottom row). As observed in living embryos (Supplementary Fig. 1b, c), descendants of anterior (Era, Ela) and posterior (Erp, Elp, Epr and Epl) ectodermal founder cells in our model became increasingly mixed with each other anterior and posterior to the IL respectively (Fig. 3c, groups I and II). In contrast, cells on either side of the IL became increasingly segregated over the analyzed time course (Fig. 3c, group III). The results of the model are consistent with the hypothesis that the IL represents a stable compartment boundary arising from the principle of energy minimization between clones of anterior and posterior cells.

**Figure 3.**
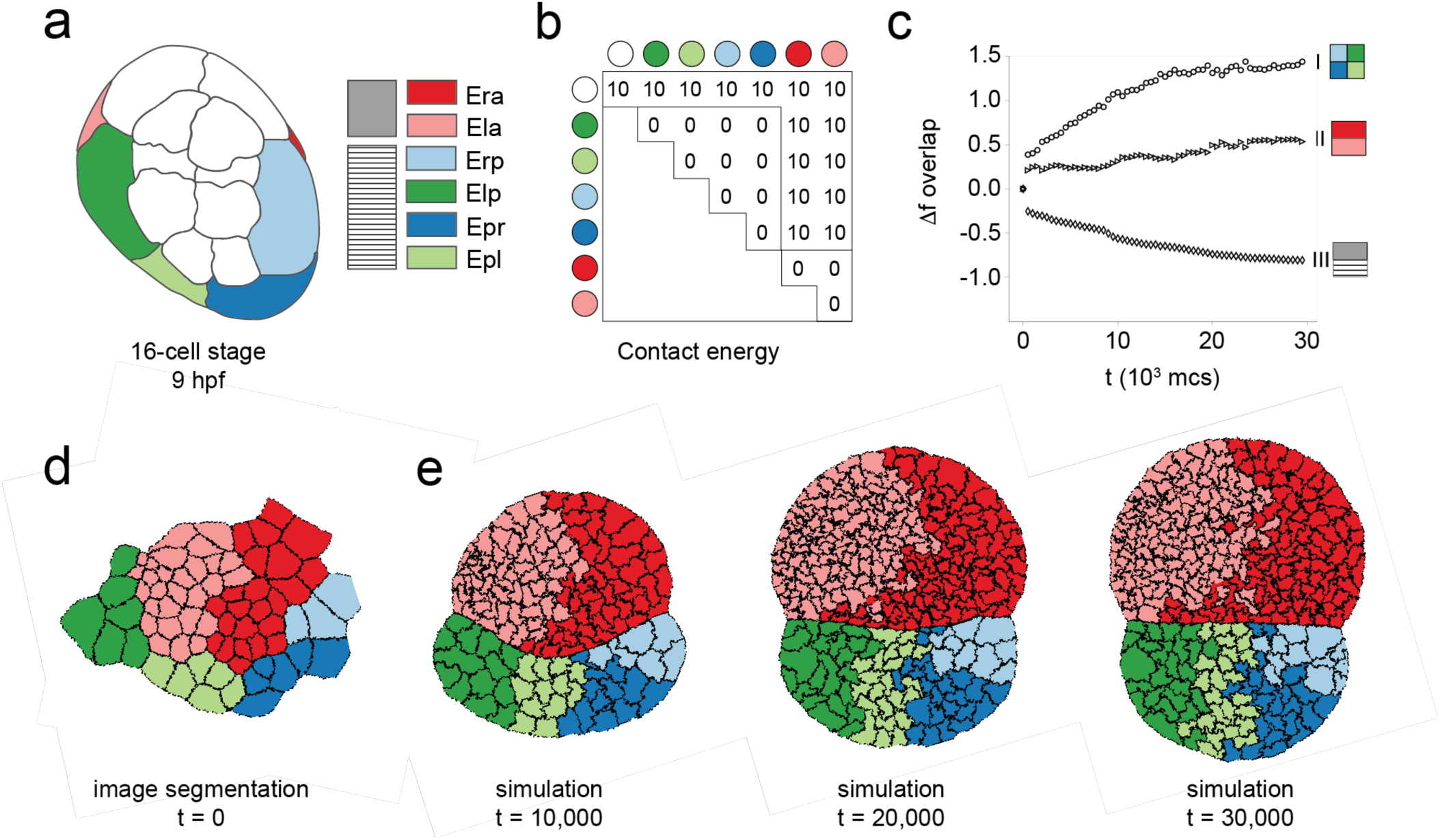
A simulation of cell sorting can explain the emergence of a straight boundary at the initiation line. **a** Tracing of a 16-cell stage embryo with the six ectodermal lineages colored. Grey and striped legend is provided to show the correspondence of these blastomere lineages to the anterior and posterior compartments shown in Figure 2. **b** Simulation contact energy inputs for each combination of lineage precursors (colored circles show blastomere identities as defined in a) as well as the environment surrounding the tissue (white circles). Higher contact energy results in decreased affinity between cells. **c** Change in normalized area of overlap plotted against time in Monte Carlo steps (mcs) for n = 10 runs of the simulation. Group I (circles) = boundaries among the four posterior lineages (Erp, Elp, Epr, Epl); group II (triangles) = boundary between the two anterior lineages (Era, Ela); group III (diamonds) = boundary between the anterior and posterior lineages. **d** Membrane segmentation of part of the conformally mapped germ disc of Embryo 1 (Supplementary Table 1) at 26 hpf. At this stage, gastrulation has already taken place, and the surface of the embryo is a continuous sheet of ectoderm. This segmentation was seeded as the starting point for the simulation. **e** Stills from three time points of the simulation of cell sorting among the 16-cell stage ectodermal lineages implemented in CompuCell3D with time denoted in arbitrary units. Cell division rates for each lineage from one of the lightsheet datasets were input into the model. Anterior is up in all embryo image panels.

### A midline of square cells forms the secondary axis of the grid through intercalation

Previous clonal analyses across multiple species of malacostracan crustaceans have shown that a morphologically distinct column of cells forms along the ventral midline of the embryo^26,27,44–48^. The cells of the midline were previously reported to be entirely comprised of Ep descendants (Supplementary Fig. 1a) in *P. hawaiensis*^27^, but are described as a polyclone descended from multiple 8-cell stage ectodermal precursors in the related amphipods *Orchestia cavimana*^46,49^ and *Gammarus pulex*^45^. We asked whether the *P. hawaiensis* midline played a role in square-cell packing along the anterior-posterior axis of the ectodermal grid, and whether its formation was lineage-dependent. We observed that *P. hawaiensis* midline formation began at 50-60 hpf through sequential mediolateral intercalation^50^ of a founder cell population (Fig. 4a-b) in a region we call the intercalation zone (IZ, Movie 4, Fig. 4a’-a”’, Supplementary Fig. 3c, d). This region subsequently split into anterior and posterior intercalation zones (Fig. 4a’’-a’”), defined as regions of intercalation anterior (IZA) and posterior (IZP) to the IL respectively, as cells aligned at the midline. The IZ founder cell population was identified in the lightsheet datasets by tracking the lineages of aligned cells at 68-70 hpf backwards to the point when only one midline cell was aligned. For the next 10 to 12 hours, the midline elongated (Fig. 4b’, c’, Supplementary Fig. 3e, f) by a combination of mitotic divisions within the founder population (Fig. 4b’, 9-cell midline) and cell intercalation. Midline intercalation started at the IL (arrowheads in Fig. 4a’-a’”, Supplementary Fig. 3c-f) and progressed both anteriorly and posteriorly, with the accumulation of midline cells in the anterior zone lagging behind that in the posterior (Fig. 4c). Cells were added to the midline sequentially (Fig. 4c’, Movie 4) and each new cell added took its final shape as a square only once it was entirely within the midline (Fig. 4b’). Once aligned, midline cells remained square and were the first square cells formed along the anterior-posterior axis of the grid (Fig. 4a’-a’”, Supplementary Fig. 3e, f). To the left and right of this axis, the remaining ectodermal cells first aligned and subsequently transitioned to square-packing in a medio-lateral wave initiated from the midline (Fig. 4a”-a’”). After the 10-12 hour period of midline elongation by intercalation and founder cell division, by 60-72 hpf the midline contained a mean of 8.75 cells in total (Fig. 3c). Subsequently, divisions of parasegment precursor rows^29^ in the posterior compartment introduced another mechanism of further midline elongation (starred cells in Fig. 4a”-a”’).

**Figure 4.**
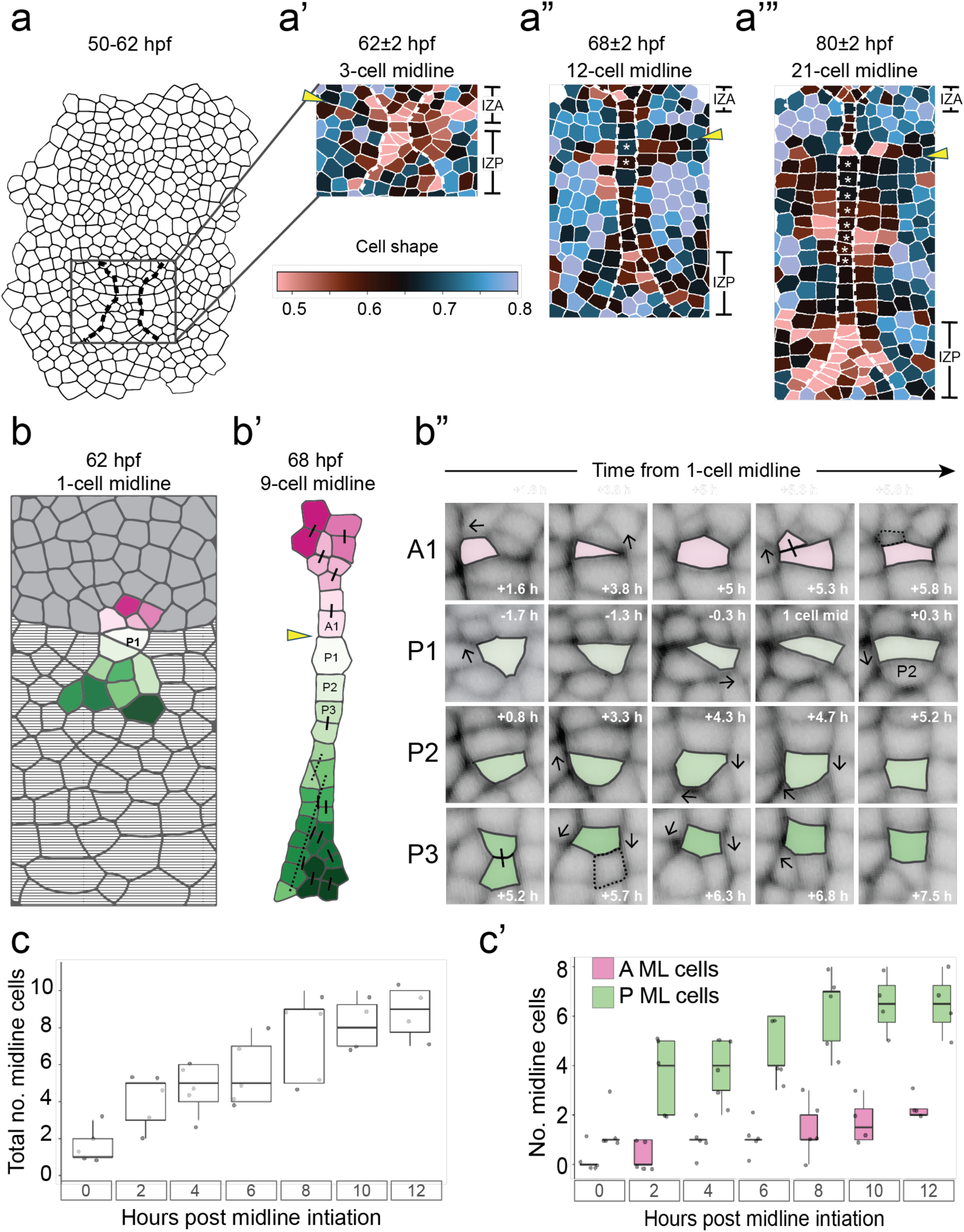
The secondary grid axis emerges as a column of square cells via the intercalation of a polyclonal founder cell population. **a** At 52-62 hpf, primarily anisotropic ectodermal cells near the ventral midline and spanning the IL (arrowheads in a’, a”, a’”), in a region we denote the intercalation zone (IZ), intercalate simultaneously in the anterior and posterior directions to form a single column of square cells forming the midline of the embryo (**a’**). **a”** The cells within the midline are the first cells to become square. Cells flanking the midline on the left and right then adopt square shapes posterior to the IL. Cells in a’, a”, and a’” are colored according to the cell shape descriptor. * in a” and a”’ indicates midline cells that have undergone parasegment precursor row duplication. **b** Cells that join the midline derive from a polyclone of founder cells that span the anterior (grey)-posterior (stripes) compartment boundary (arrowhead). **b’** Founder cells intercalate in both anterior (IZA) and posterior (IZP) intercalation zones (as shown in a’, a” and a’”), adding or subtracting sides until they form a square. Arrows indicate the direction of a growing or shrinking cell side. Dotted outlines in rightmost panel of top row and second panel of bottom row indicate the outlines of the sister cell resulting from the division in the previous panel. **c** Numbers of total cells in the midline over time (n = 5 embryos). Midline initiation is defined as the time when the first posterior cell (P1) is aligned. By 12 hours after midline initiation, a mean of nine total cells have been added to the midline. **c’** Numbers of cells in the midline anterior (A ML: pink) and posterior (P ML: green) to the IL over time (in = 5 embryos). Anterior is up in a-a”’ and b-b’’.

In contrast with the initiation line, the midline grid axis did not strictly correspond to clonal boundaries between 16-cell ectodermal lineage domains (Supplementary Fig. 3). The emergence of the midline, with the same cell behaviors extending it in both directions along the anterior-posterior axis, spanned the anterior and posterior lineage domains (Fig. 4b, Supplementary Fig. 3c, d). However, we observed that the posterior midline was not exclusively made up of Ep descendants (Supplementary Fig. 3c). This is in disagreement with the original lineage tracing report for *P. hawaiensis*^27^, but is consistent with the report that embryos lacking an Ep blastomere still form a midline^40^. In the anterior portion of the midline, clones from both anterior ectodermal lineages (Era and Ela) contributed to the midline founder population with no apparent relationship between lineage domains and the anterior IZ (Supplementary Fig. 3a, b). Taken together, these results suggest that the midline initiates the emergence of complete square cells and square-cell packing along the anterior-posterior axis through a highly coordinated morphogenetic process based on cell recruitment rather than inherited fates, as it is not dependent on the 16-cell stage ectoderm lineages (Supplementary Fig. 3).

### The initiation line and midline column form tensile cytoskeletal cables

Given the emergence of square-cell packing geometry first at the IL (around 30 hpf) and subsequently at the midline (around 50 hpf), we hypothesized that these boundaries introduce tension into the tissue to initiate the transition to square-packing. To test this hypothesis, in *P. hawaiensis* embryos stained with phalloidin (Fig. 5a) and injected with a myosin 2 mRNA reporter construct (based on the *P. hawaiensis* ortholog of the *D. melanogaster* gene encoding the regulatory light chain of the non-muscle type 2 myosin, known as *spaghetti-squash* ^51^ (*Ph-sqh*), Supplementary Fig. 4), we measured the relative amounts of filamentous actin (F-actin) and myosin 2 at the IL when it appeared in earlier stages (38-56 hpf, Fig. 5a-a’), and on either side of the midline when it formed at later stages (around 68-74 hpf, Fig. 5b, b’).

**Figure 5.**
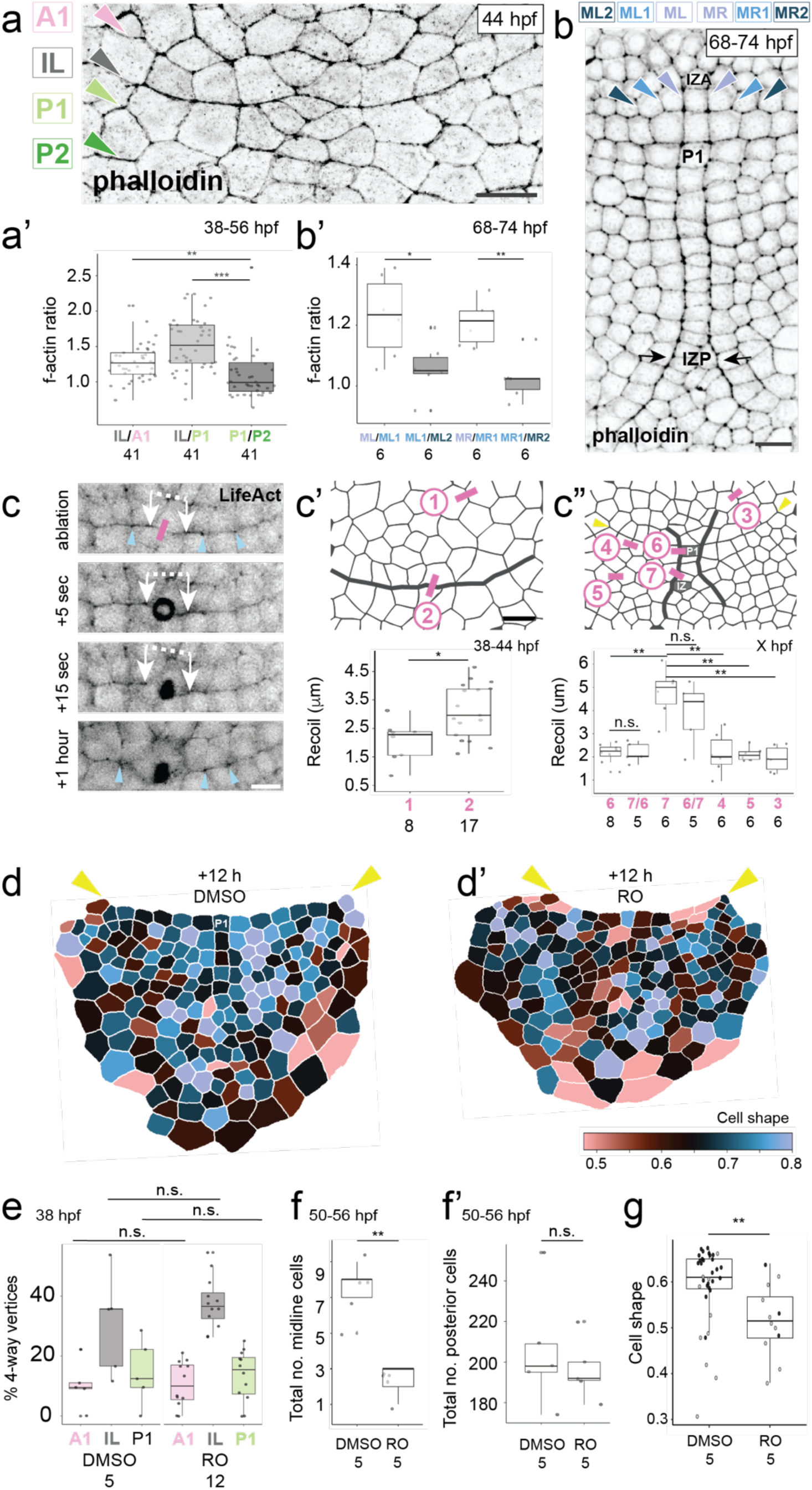
The initiation line and midline borders are tensile cytoskeletal cables. **a** Maximum intensity projection of 20 0.5µm optical sections of the ventral ectoderm of a 44±2 hpf embryo stained with phalloidin to label F-actin. **a’** Ratios of mean F-actin intensity measured in 38-56 hpf embryos. The regions quantified (A1, IL, P1, P2) are shown in **a**. Each region begins at the arrowhead on the right of the micrograph and extends across approximately 10 contiguous cell sides to the left side of the micrograph. **b** Maximum intensity projection of six 0.9µm optical sections of a 68-74 hpf embryo with a 15-cell midline; F-actin is labeled with phalloidin as in a. Arrowheads indicate the six regions (lines running parallel to the anterior-posterior axis, named MR, MR1, MR2, ML, ML1, ML2) where F-actin intensity was quantified. Each region quantified begins at the arrowhead shown in **b** and extends towards the posterior along contiguous cell sides to the posterior IZ (black arrows). **b’** Ratios of mean F-actin intensity measured in 68-74 hpf embryos with 7-15-cells in the midline. **c** 38-44 hpf embryo subject to laser ablation at the IL. **c** Pink rectangle indicates the ablation site; white arrows connected by a dashed line indicate the lengths measured to calculate recoil; blue arrowheads at IL indicate vertices surrounding the incision site. **c’** Ablation sites (top: thick line indicates the IL) and recoil plot (bottom) in 38-44 hpf embryos in which recoil was measured following IL ablation. **c”** Ablation sites (top: thick lines indicate the cables demarcating the nascent midline) and recoil plot (bottom) in 58-69 hpf embryos with 3-7 cells in the midline. Pink numbers on X-axis categories indicate the ablation site as shown in the schematic above. 7/6 indicates that a first incision was made at position 6 (next to the midline P1 cell/at the compartment boundary), followed by a second incision at position 7 (posterior IZ). In 6/7, the same two incisions were made in reverse order. **d** 1-cell midline embryos (50-56 hpf) were incubated in 50µM DMSO or Rockout (**d’**), and the cells of the posterior compartment were segmented and their cell shape descriptor was color-mapped 12 hours later. The IL is indicated by yellow arrowheads. **e** Percentage of 4-way vertices in 38 hpf embryos incubated in DMSO and 50µM Rockout for 12 hours. P-values for comparisons between DMSO and RO-treated samples as follows: A1: 0.48; IL: 0.18; P1: 0.48. **f** Total number of midline cells is significantly decreased in 62-68 hpf embryos treated with Rockout for 12 hours (p-value = 0.004). **f’** Total number of cells in the posterior compartment of 62-68 hpf embryos treated with Rockout for 12 hours is not significantly different from DMSO controls (p-value = 0.35). **g** Cell shape values for cells aligned into the midline in 12-hour DMSO (n = 35 cells) and Rockout (n = 12 cells) treated embryos. Grey symbols indicate the single cells at the anterior-most and posterior-most positions of the midline; black symbols indicate all other midline cells. Samples are the same as those plotted in f and f’ (p-value = 0.002). * p-value < 0.05; ** p-value < 0.01; *** p-value < 0.001; n.s. not significant (Mann Whitney U-test; all tests one-tailed). Scale bars for a, c’, d = 25µm and c = 15µm. Anterior is up in all micrographs. Sample sizes (number of embryos) in all plots indicated by numbers underneath category labels.

In 38-56 hpf embryos, we found significantly higher levels of F-actin (Fig. 5a, a’) and myosin 2 (Supplementary Fig. 4a, a’) at the IL relative to the equivalent region of interest (ROI) one cell anterior (IL/A1) or posterior (IL/P1), compared with one to two cells posterior (P1/P2).

Similarly, we also found significantly higher levels of F-actin on the left (ML) and right (MR) sides of the midline in embryos with 6-15 cells in the midline (68-74 hpf) (Fig. 5b, b’) compared with the equivalent ROIs one and two cells to the left (ML1, ML2) and right (MR1, MR2). In contrast, levels of myosin 2 on either side of the midline were not significantly higher than the ROIs to the left and right of the midline (Supplementary Fig. 4b’b”). We hypothesize that these accumulations of cytoskeletal proteins constitute supracellular cables running along the two axes of the ectodermal grid (Supplementary Fig. 5).

In *D. melanogaster*, actomyosin cables segment embryonic and larval tissues by introducing local tension into the tissue^19,52,53^. We hypothesized that the supracellular cables we detected in the *P. hawaiensis* germ band were similarly under tension, allowing them to regulate the formation of the first square cells in the grid. To test this hypothesis, we cut the cables with a pulsed laser tuned to 927 μm and measured subsequent tissue recoil around the incision site (Fig. 5c, dashed line and arrows). In 38-44 hpf embryos that had formed the IL, recoil speeds were significantly (1.5 to 2.5-fold) higher at the IL compared with control cuts made four to seven cells anterior to the IL (Fig. 4c’). Interestingly, this increase is comparable to the mean of 2.37-fold higher tension reported at the *D. melanogaster* wing disc anterior-posterior compartment boundary compared with that at cell edges within the anterior or posterior tissue compartments^52^. In embryos live-imaged following the IL cut, we observed that the 4-way vertices surrounding the incision site remained intact (black arrowheads in Fig. 5c), suggesting that maintaining a square shape at the IL is not dependent on tension in neighboring cells.

In later stage embryos in which 3-6 cells had aligned into the midline (58-69 hpf), recoil at midline cables in the posterior IZ was significantly (2 to 3-fold) higher than at other non-IZ regions one to four cell diameters anterior or lateral to the midline (Fig. 4c”). In contrast, recoil speeds at the cables in the anterior region of the midline where intercalation had finished and cells were square in shape (P1 in Fig. 5b, c”) were not significantly different than those at the lateral and anterior control regions (Fig. 5c”). To determine whether tension was mechanically coupled across cells contributing to the cable, we measured the recoil of a second cut in the midline after making a first cut two to three cells away in the cable (Fig. 4c’), and found that P1 and IZ second cut recoils were not significantly different from those at first cuts (Fig. 5c”). We note that a similar phenomenon of local, potentially cell-autonomous contribution to tension across a supracellular actomyosin cable spanning many cell diameters, has also been reported in the morphogenesis of an extra-embryonic membrane in the beetle *Tribolium castaneum*^54^. Taken together, these results suggest that there are supracellular cables under tension at the IL and at the midline, and that tension within midline cables is heterogeneous and locally generated.

Tension in actomyosin cytoskeletal cables results from the phosphorylation of the regulatory light chain of myosin 2, which slides the attached actin filaments past one another^55^. The results of our laser ablation experiments suggested that while both the IL and the midline cables possessed supracellular actomyosin cables that were under tension, tension at IL was not essential for the formation of 4-way vertices. This led us to hypothesize that abrogation of myosin 2 phosphorylation would not hinder square cell-packing geometry formation at the IL at earlier stages, but might disrupt square cell formation at the midline at later stages.

To test this prediction, we first assessed the effect of sustained elimination of myosin 2 phosphorylation during the later stages. We focused on the period when intercalation is the only mechanism of midline formation (50-68 hpf, Fig. 4b, b’), and chose morphologically stage-matched embryos in which just one cell had aligned into the midline (50-56 hpf, Fig. 4b) for incubation in 50µM of the Rho-kinase inhibitor Rockout for 12 hours (Fig. 5d, d’). This concentration of Rockout did not impact normal cell proliferation (Supplementary Fig. 6a, b), thereby allowing the thresholds of contractility needed for cytokinesis^19,53^. Rockout-treated embryos at this stage had significantly fewer cells aligned into the midline than controls (mean 1.4 vs 6.4, Fig. 5f) but maintained the same total number of cells in the posterior compartment (Fig. 5f’). Notably, a relatively straight boundary at the IL was maintained in the midline Rockout-treated embryos (Fig. 5d’, arrowheads). In contrast, earlier stage embryos (38 hpf) treated for 12 hours in 50µM Rockout had, by 50 hpf, percentages of 4-way vertices that were not significantly different from controls (Fig. 5e). Taken together, these results suggest that while 4-way vertices first appear at the IL (Fig. 2), their maintenance is not dependent on the phosphorylation of myosin 2 (Fig. 5e). In contrast, however, subsequent establishment of the first square cells is dependent on the contractility of myosin 2 within midline cables (Fig. 5g).

### The midline cables organize square-cell packing and the emergence of a stripe of Engrailed-positive cells

Given the presence of differential tension within the germ disc (Fig. 5c, c”), and the midline as the initiation site of the transition to square-packing along the axis of grid elongation (Fig, 4, Supplementary Fig. 3), we hypothesized that localized mechanical perturbation of the midline would disrupt the organization of square-cell packing across the tissue, and that perturbing square-cell packing would in turn have consequences for the correct establishment of segments. To test this, we performed two classes of physical ablations, either directly at the midline (Mid), or three cell diameters lateral to the midline (Lat) in embryos in which 3-6 midline cells had aligned (58-69 hpf, A: pink rectangles in Fig. 6b, c). The contralateral side of the same embryo, which was not ablated (NA: pink rectangles in Fig. 6b, c), served as an internal control. In the resulting four experimental groups (Mid-A, Mid-NA, Lat-A, Lat-NA), we analyzed posterior compartment cell shapes (Fig. 6d, d’), number of cells in the midline (Supplementary Fig. 7), and expression of the segment marker Engrailed^37^ (En, Fig. 6e, Supplementary Table 3) 9-12 hours after ablation. In the 100 closest cells to P1 posterior to the IL, we used Kernel density estimation^56,57^ to estimate the cell shape probability densities for all groups, including wild type embryos morphologically stage-matched to Lat embryos based on the number of cells in the midline. We plotted the deviation of the four experimental groups from the wild type distribution over the range of cell shape values observed (Fig. 6d’).

**Figure 6.**
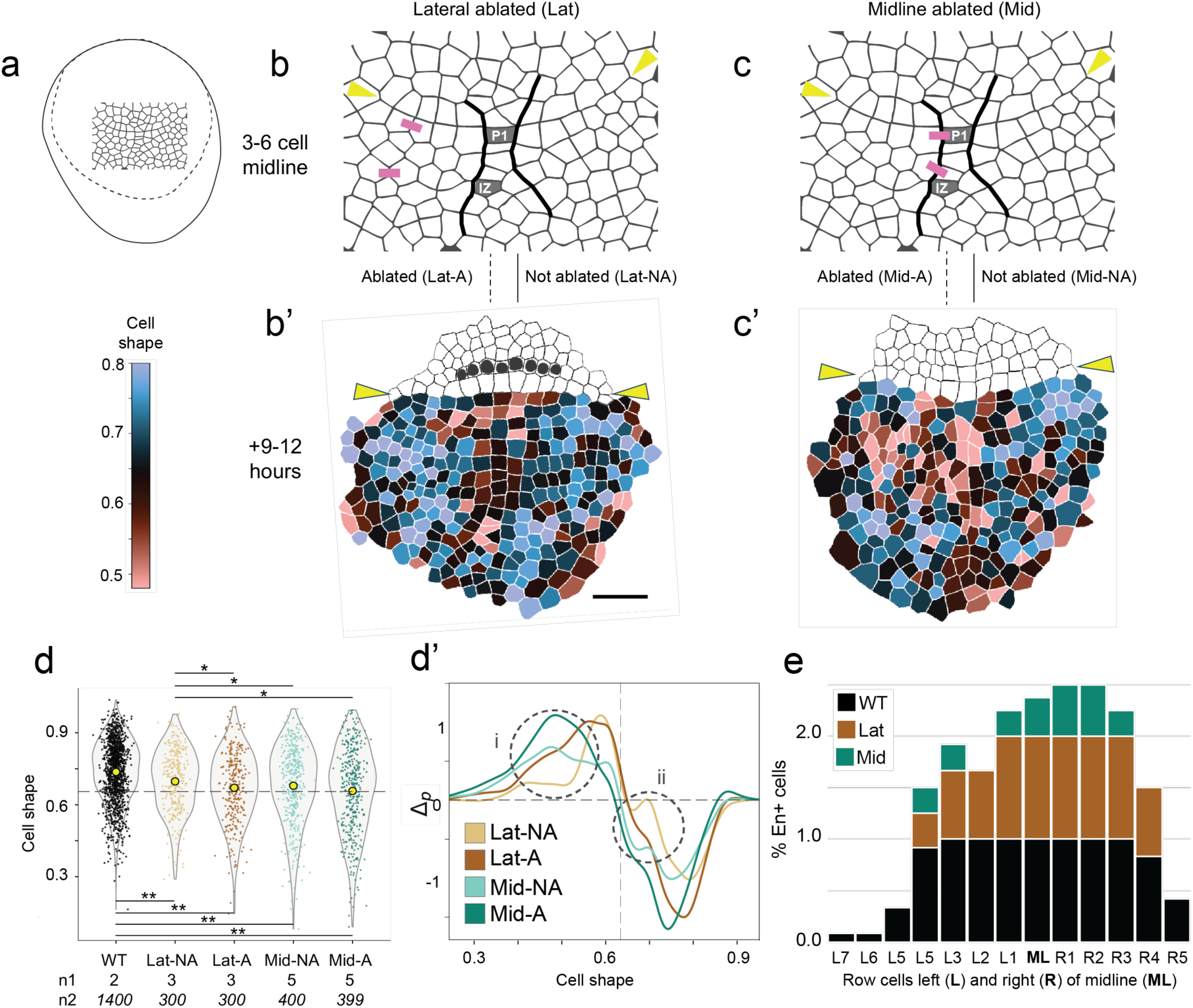
The midline cables organize square-cell packing and the emergence of the first complete stripe of Engrailed-positive nuclei. **a** The region within the germ disc where ablations were made, delineated by the dotted line. **b, c** Locations of laser ablations performed in different regions of the ventral epithelium in 58-69 hpf embryos with 3 – 6 cells in the midline. **b’, c’** Cell shape descriptor values and En expression 9-12 hours after ablation. **d** Cell shape descriptor value distributions across treatments, including unablated embryos (WT) of equivalent chronological age. Yellow circles indicate the mean, dotted line the value for an isometric square. Only significantly different pairwise comparisons (based on the bootstrapping approach described in Methods) are indicated as follows: * p < 0.05; ** p < 0.01. Sample sizes indicated by n1: number of embryos and n2: number of cells. **d’** Deviation of the four ablated groups from the WT probability density (Δ_p_ = 0) over the range of cell shape values observed. A positive value indicates shapes gained and a negative value indicates shapes lost. Dotted vertical line indicates the value for an isometric square; colors are the same as in panel d. **e** Proportion of Mn cells (categories on horizontal axis) expressing En in wild type and in embryos subjected to lateral (brown) or midline (green) ablations shown in b and c respectively. Data in Supplemental Table 3.

We found that at 38 hpf, ablations of the IL and control ablations three cell diameters anterior to the IL did not prevent the formation of square cells nor the establishment of the ectodermal square grid (Supplementary Fig. 8). Additionally, 38 hpf embryos in which we made a single cut in the IL cable maintained a straight line at the IL region of the tissue and began midline formation 12 hours later (Supplementary Fig. 8). This is consistent with the observation that 4-way vertices still form correctly after inhibition of myosin 2 phosphorylation at IL establishment stages (Fig. 5e).

In contrast, in embryos in which 3-6 midline cells had aligned (58-69 hpf), ablations lateral to the midline significantly perturbed the cell shape distribution on the ablated side of the embryo (Lat-A, Fig. 6d) compared with the unablated side (Lat-NA), overall driving them towards a rounder, rather than square, shape (Fig. 6b’, d’). Relative to wild type, Lat-NA exhibited the smallest increase in anisotropic cells (defined as those with a cell shape metric value of ⩽0.5; circle (i) in Fig. 6d’) and the smallest decrease in square cells (circle (ii) in Fig. 6d’) of all four groups.

Unlike the localized impact of lateral ablations, midline ablations significantly disrupted cell shape distributions both on the ablated side (Mid-A) and on the contralateral control side (Mid-NA) (Fig. 6c’,d’). On both sides of midline ablated embryos, more cells took on more irregular shapes, characterized by lower values of the cell shape metric (Fig. 6d). In particular, the proportion of square cells was lower than wild type in both the experimental and the control (contralateral) group of midline ablations (circle (ii) in Fig. 6d’). Midline ablation destroyed the organization of the square-packed grid throughout the posterior germ rudiment (Fig. 6c’). In sum, the effect of the lateral ablation was minimal and localized to the region immediately around the ablation, while perturbing just one of the midline cables had a tissue-wide impact on cell shapes, disrupting square cell formation and abolishing the square ectodermal grid.

Midline, but not lateral, ablations also disrupted midline elongation. Laterally-ablated embryos developed midlines with total numbers of cells comparable to wild type (12-19 cells, Supplementary Fig. 7) while midlines in midline-ablated embryos were shorter (5-10 cells, Supplementary Fig. 7). This suggests that midline cable ablations, but not ablations lateral to the midline, disrupted posterior intercalation at the midline. These data are consistent with the hypothesis that the alignment of cells into the midline, driven by the contractility of the midline cables, organizes cell shape across the tissue.

Finally, we asked whether establishment or maintenance of the square cell grid was important for correct (para)segment patterning. At stage 11-12 of embryogenesis (66-68 hpf^29^), when midline elongation by intercalation has completed (Fig. 4a”), the protein product of the segment polarity gene *engrailed* (En) becomes detectable in two cells anterior to the IL, initially in a few cells within the same row at or near the midline^29^. The En expression domain subsequently expands in an increasing, left-right symmetrical number of lateral cells within the row^29^. This expression marks the prospective mandibular segment (Mn, arrowhead in Fig. 1i, j)^29,44^. Subsequent rows of En-expressing ectodermal cells, delineating the remaining segments of the body plan, arise correlated with the same stereotyped cell divisions of the cells in the grid that contribute to midline elongation (Fig. 1j, k) ^29,44^.

Given that En expression in the prospective mandibular segment is detected right after the midline intercalation phase that appears to create the first square cells in the grid, we asked whether this expression was dependent on the tensile midline cables in the same way as square cells and square-packing order. Across the WT and ablated samples, we found that the total number of Engrailed-positive nuclei in the prospective mandibular segment was positively correlated with the number of aligned midline cells (Supplementary Fig. 7b, Supplementary Table 3). In other words, as the midline became longer, more lateral cells within Mn expressed En. Strikingly, however, midline ablation reduced the number of En-positive Mn nuclei, even on the contralateral control side of the embryo (Figure 6e, Supplementary Table 3). In contrast, lateral ablations had little impact on the number of Engrailed-positive Mn nuclei on either the experimental or the control side of the embryo (Figure 6e, Supplementary Table 3). Midline but not lateral ablations also decreased the regularity with which Mn cells expressed En (Figure 6e, Supplementary Table 3). Given these results, we hypothesize that the first establishment of En expression across a complete segment, as seen in Mn, requires midline-cable driven tissue organization.

## Discussion

### The role of cell lineage in initiating square cell formation

We found that descendants of the 16-cell stage ectodermal founder blastomeres segregate into anterior and posterior compartments, and that the resulting compartment boundary (IL) forms the first axis of the square cell grid. This suggests the possibility that cell lineage at least partly underlies square cell morphogenesis. The results of our simulation suggest that cell sorting between the cells of the anterior and posterior compartments, a phenomenon observed in multiple contexts in animal development including amphibian embryos^58^ and *Drosophila* imaginal discs^59^, regulates the formation of a straight compartment boundary. The mechanism that regulates this cell sorting could in principle be one or more of differential cortical stiffness^60–62^, differential adhesion^61–67^, or a mechanical barrier imposed by the supracellular actomyosin cable that forms at the IL. Relevant to the latter hypothesis, we note that although the cable at the IL is under tension, coupled tension and the phosphorylation of myosin 2 are not required to maintain 4-way vertices at this boundary. This suggests that local tension at this IL cable is not necessary to maintain the compartment boundary. Alternatively, it is possible that the interaction between the anterior and posterior lineages can restore the tensile cable upon ablation.

The results of previous experiments demonstrating that the *P. hawaiensis* early embryonic ectoderm has regulative capacity are interpretable as consistent with the cell sorting hypothesis^40^. In these experiments, individual 8-cell stage ectodermal blastomeres were photoablated and the remaining two blastomeres were injected with a lineage tracer. The results showed that El and Er can replace each other’s anterior descendants, while Ep can replace their posterior descendants^40^. We hypothesize that compensation within the ectoderm, and ultimately grid formation, is possible only when progenitors of both anterior and posterior lineages remain, and that cells from both compartments are necessary to initiate square cell-packing geometry at the IL. Under this hypothesis, both anterior and posterior cell types would be needed to form the boundary because each lineage would contribute a distinct molecular state (e.g. cortical stiffness and/or adhesive properties), the combination of which would then lead to emergent cell sorting.

In embryos of diverse animal species–including *D. melanogaster*^68–70^, the flour beetle *Tribolium casteneum*^54^, the cnidarian *Hydra vulgaris*^71^, the zebrafish *Danio rerio*^72^, and the quail *Coturnix japonica*^73^–mechanically integrated actomyosin cables spanning multiple cells play crucial roles in epithelial morphogenesis. We hypothesize that differential genetically-determined fate between the anterior and posterior lineages also underlies the assembly, and potentially the shape organizing function, of the cytoskeletal cable we observed at the anterior-posterior boundary (IL) from 38 hpf onwards, similar to anterior-posterior boundary formation in the *D. melanogaster* wing disc^52^. In the early *Caenorhabditis elegans* embryo, both endodermal cell fate and the phosphorylation of myosin 2, which are necessary for gastrulation in endodermal cells, are dependent on the Wnt-Frizzled signaling pathway, suggesting a direct connection between cell fate and morphogenesis^74^. A similar relationship could exist within the *P. hawaiensis* ectoderm, the molecular determinants of which have yet to be identified.

Further ablation experiments in 16-cell stage embryos that eliminate all anterior or posterior lineages, or elucidation of the transcriptomic, proteomic and mechanical states of the anterior and posterior compartment clones, would be useful to elucidate the mechanism of cell sorting at the IL. A hypothesis of one or both of differential adhesion or differential cortical stiffness would predict that anterior and posterior cells have different levels of cell-cell adhesion molecules and/or different cortical cytoskeletal behaviors. The fact that the IL marks a clonal boundary could in principle provide a heritable mechanism of achieving differential molecular states between clones. For example, anterior cells might inherit a level of adhesion molecules or a cortical cytoskeletal conformation distinct from that of their posterior neighbors by virtue of differential inheritance of cytoplasmic factors, or through inheritance of a gene regulatory state.

### Lineage dependent and lineage-independent mechanical cues establish square cell packing

Once the IL has straightened into an actomyosin rich boundary axis, we found that a midline cell founder population begins to intercalate perpendicular to the IL such that square cells are added sequentially to the midline between two actomyosin cables. This grid axis does not correlate with 16-cell lineage domain boundaries, such that the midline founder population is a polyclone containing descendants of multiple ectodermal founders (Supplementary Fig. 1, 2). These findings suggest that, unlike the IL, the formation of the midline and its cables is not regulated by inherited differences among these lineages. We hypothesize that the midline grid axis is instead initiated by mechanical stimuli, similar to how the formation of the primitive streak in early avian embryos, which involves anterior-posterior oriented actomyosin cables^75^, is driven by tissue flows that regulate the expression of fate-determining genes.^9^ Under this hypothesis, the midline founder population would become defined–and, in turn, potentially genetically differentiated–as a result of tissue-wide forces. Future work could examine whether fate-determining genes of this population include components of the planar cell polarity signaling pathway, which is involved in medio-lateral intercalation of cells in multiple developmental contexts.^76–80^

We have provided evidence that the mechanical integrity of these midline cables is necessary for the formation of isometric cells, the alignment of cells into rows, the maintenance and packing of square cells across the ectodermal grid, and the establishment of segmentally iterated En expression. We hypothesize that these cables drive tissue-wide mechanics, just as tissue flows pattern the retina^81^ and embryonic germ band extension^82^ in *D. melanogaster*, or how supracellular actin cables organize whole body regeneration in *H. vulgaris*^71^. In this framework, the tension introduced by the midline cables at the intercalation zones could be the initial condition necessary to trigger and perhaps subsequently balance the transition to square-packing. The variance in recoil times in response to ablation at the IL and midline intercalation zones (Fig. 5c’, c”), suggests that tension at these regions could be dynamically modulated in response to global tissue forces.

### The relationship between square-cell packing, gene expression, and segmental patterning

In ascidian and mouse embryos, cell geometry is crucial for the expression of fate-determining genes^83,84^. Our study suggests that in *P. hawaiensis*, square grid morphogenesis and subsequent segment formation emerge from axes that are established according to differing degrees of instruction from tissue mechanics and genes. We also provide evidence that expression of the segment polarity gene *en* in the *P. hawaiensis* ectoderm is sensitive to mechanical perturbation and dependent on the formation of the midline. Reminiscent of this finding, in *D. melanogaster* embryos injected with paramagnetic-beads and subjected to ectopic tissue-wide forces with magnetic tweezers, expression of En at parasegment boundaries was differentially altered according to the location and magnitude of the force^85^. The authors of that study observed a change in nuclear shape and redistribution of myosin 2 in tissue regions experiencing the applied force, suggesting that expression of *en* is downstream of these mechanical and geometric parameters^85^. This raises the possibility that mechanical regulation of *en* expression could be a conserved feature in multiple pancrustacean species.

In early *D. melanogaster* embryos, membrane fluidization by an ether treatment applied to whole embryos results in larvae with a dramatic range of segmentation defects, from complete loss of segments to the elimination of only a single segment, which resembles segmentation gene loss of function phenotypes^86^. Our *P. hawaiensis* midline ablation experiments, in which cuts were made in the exact same cell positions across samples, similarly resulted in a range of changes in cell shape, midline cell number, and *en* expression, suggesting that segmentation in *P. hawaiensis* could arise from a balance of tuned forces within the ectodermal epithelium. In this mechanical framework, the ultimate effect of midline cable disruption could depend on the mechanical state elsewhere in the tissue. Further experiments determining the longer-term effect of early midline disruption, and the alteration of square-cell packing, on the formation of more posterior segments will be necessary to establish the extent of the midline cables’ organizing capabilities.

Beyond amphipods, previous comparative studies of embryogenesis in other malacostracan orders (including Cumacea, Tanaidacea, Isopoda, and Mysidacea) have found that they generate some of the cell material for their ectodermal grids through consecutive divisions of ectodermal stem cells (ectoteloblasts) that provisionally align their clonal progeny into rows in the posterior portion of the grid^87^. However, ectodermal grids in these species also contain cells derived from non-ectoteloblast lineages, suggesting that some other mechanism of tissue organization must be present^28^. Amphipod grids like that of *P. hawaiensis* completely lack ectoteloblasts and thus must be organized by what Wolfgang Dohle and Gerhard Scholtz called a “row-forming factor,” which they hypothesized to be a “matrix forcing the cells into a grid-like pattern”^87^. Our results suggest that this “matrix” emerges first from tensile cell-sorting at the IL and subsequently from cytoskeletal cables at the midline. *O. cavimana*, another amphipod, exhibits stereotypic early cleavage patterns similar to that of *P. hawaiensis*, suggesting that fate determination within the 8-16 cell embryo could also play a role in grid morphogenesis of this species^46^. Further lineage studies across the Amphipoda, the early embryos of which all undergo holoblastic cleavage^88^, are needed to understand the full extent of this mechanism within this group.

The amphipod anterior-posterior compartment boundary corresponds to a boundary classically called the naupliar-post-naupliar boundary. Under this concept, the naupliar region of the germ band, is the anterior region that will give rise to the head and mandibular segments, formed through a combination of relatively uncoordinated mitoses, cell intercalation, and cell rearrangements, and the post-naupliar region. In contrast, the post-naupliar germ band is formed through the activity of either ectoteloblasts or segment precursor row duplications^44,45,89^, and gives rise to the trunk segments and telson. Interestingly, our work shows that the IL, positioned one cell diameter posterior to the first complete segmental stripe of En expression in the prospective mandibular segment, corresponds to the classically defined naupliar-post-naupliar boundary.

The anterior-posterior compartment boundary is established by the descendants of ectodermal lineage precursors defined by stereotypical holoblastic cleavages^27,33^ and likely also by asymmetric inheritance of lineage potential to early blastomeres that begins at first cleavage^35,90,91^. However, holoblastic cleavage is a derived state for the Malacostraca, in which many extant species undergo syncytial cleavage^46^, and the ancestral cleavage state is thought to be meroblastic (in which cleavage furrows form but cytokinesis is not completed, effectively resulting in a syncytium but with some partial yolk partitioning by the formation of incomplete furrows)^92,93^. We speculate that the evolution of holoblastic cleavage in amphipods facilitated the emergence of lineage-based IL formation and the anterior-posterior compartment boundary as a mechanism for establishing the ectodermal grid and generating the post-naupliar germ band. Under this hypothesis, the evolution of an anterior-posterior boundary among the ectoderm clones eliminated the need for patterning introduced by a row of proliferating ectoteloblasts within the germ disc. With regards to grid organization along the anterior-posterior axis, we hypothesize that midline formation via intercalation, and the importance of the mechanical organizing properties of the midline for establishing correct segmental patterning, are likely to be conserved across the Malacostraca. Further comparative studies should determine whether midline cables are present in other species.

## Methods

### Animal culture and transgenesis

*P. hawaiensis* were cultured at 26 °C in 16-quart plastic tanks containing 1.022 specific gravity Instant Ocean artificial sea water (hereafter ASW) made with distilled water and fed a diet of TetraPro PlecoWafers and fresh carrots. For live imaging of nuclei (Fig. 4c) we used a previously reported^94^ transgenic *P. hawaiensis* line in which the *D. melanogaster* histone H2B fused with a monomeric Red Fluorescent Protein is under control of a *P. hawaiensis* heat-shock promoter (hereafter *H2B-mRFPruby*).

### Embryo collection and microinjection

Embryos for all experiments were collected and microinjected according to established protocols^95^. Embryos were collected into and cultured in ASW filtered with a 0.22µm filter (hereafter FASW). For precise staging and stage-matching among broods, only 1-cell through 8-cell stage embryos were collected. All microinjected embryos were injected at the late 1-cell stage once the chorion had hardened. Microinjected embryo cultures were supplemented with 1:100 Penicillin-Streptomycin (10,000 U/mL penicillin, 10,000 µg/ml streptomycin; Gibco 15140-122) and 1:200 Amphotericin-B (250 µg/ml, Gibco 15290-026). The culture medium was changed out every 1-2 days. For live imaging of the actomyosin cytoskeleton, embryos were injected with 180pl of 2 µg/µl of mRNA reporter constructs encoding either the actin-binding peptide LifeAct^96^ fused to the monomeric mNeonGreen fluorescent protein^97^ (hereafter *LifeAct-mNeonGreen* or *LA-mNG*) or the *P. hawaiensis* myosin II regulatory light chain gene (*Ph-sqh*) fused to mNeonGreen (hereafter *Ph-sqh-mNG*). In vitro transcribed capped mRNAs encoding these reporters were synthesized with the mMessage mMachine T7 kit (Invitrogen AM1344) from plasmid templates pBlueSK-LifeAct-mNG and pBlueSK-Phsqh-mNG linearized with the restriction enzyme NotI, as previously described (Kontarakis 2014). Plasmid templates pBlueSK-LifeAct-mNG and pBlueSK-Phsqh-mNG were generated by PCR amplification of the *LA-mNG* and the Ph-sqh-mNG open reading frames (with primer combinations pBlue_LA_GBS_F 5’-agaactctgggggatctgatAatgGGCGTGGCCGATC-3’ / pBlue_mNG_GBS_R 5’-cggttcagaacttataacaTTACTTGTACAGCTCGTCCATG-3’ or pBlue_Phsqh_GBS_F 5’-agaactctgggggatctgatAATGTCGTCCCGCAAAGC-3’ / pBlue_mNG_GBS_R) from vectors pSL-PhHS-LA-mNG and pSL-PhHS-Phsqh-mNG, respectively (Kalogeridi and Pavlopoulous, personal communication). PCR amplicons were cloned with Gibson assembly into ClaI-digested pBlueSKMimRNA^98^, replacing the Minos transposase ORF with the LA-mNG or the Ph-sqh-mNG ORFs, respectively. For long-term lightsheet imaging of cell membranes and nuclei, embryos were co-injected with in vitro transcribed capped mRNAs synthesized with the mMessage mMachine SP6 kit (Invitrogen AM1340) either from plasmid template pEXPTol2-H2A-mCherry (encoding the nuclear fluorescent reporter H2A-mCherry) linearized with the restriction enzyme BglII^99^ or from plasmid template PCS2 + Lyn-EGFP (encoding the membrane fluorescent reporter Lyn-EGFP) linearized with the restriction enzyme NotI^100^.

### Immunohistochemistry and drug treatments

Embryos were dissected away from their outer membrane according to established protocols^101^ and fixed for one hour at room temperature in 4% paraformaldehyde made up in FASW. Fixed tissue was immediately transferred into a blocking solution (1x phosphate buffered saline (1x PBS) + 5% normal goat serum + 0.1% Triton X-100) or stored at 4 °C in 1x PBS + 0.1% Triton X-100 for up to three days prior to staining. Embryos were stained with 1:100 AlexaFluor 488 Phalloidin (Invitrogen A12379) or Rhodamine Phalloidin (Invitrogen R415) for F-actin, 1:1000 rabbit anti-Myosin 2 656^102^, 1:80 mouse anti-Engrailed Mab 4D9^37^, 1:500 mouse anti-phospho-histone H3 (Cell Signaling 9706S), or 1:50 rabbit anti-cleaved caspase-3 conjugated to AlexaFluor 555 (Cell Signaling 9604S) overnight at 4°C. We followed these incubations with a solution containing 1:10,000 Hoechst 33342 (Sigma B2261, 10 μg/μl stock solution) as well as secondary antibodies 1:500 AlexaFluor 647-conjugated Goat Anti-Mouse (Invitrogen A31625) or 1:500 AlexaFluor 555-conjugated Goat Anti-Mouse (Invitrogen A31621) in blocking solution for two hours at room temperature. Prior to fixation, drug-treated embryos were cultured for 12 hours at 26°C in a solution of Rockout (Rho Kinase Inhibitor III, Sigma 55555-3) dissolved in anhydrous DMSO and added to a 30mm petri dish containing 6ml of 50% FASW, as previously described^36^. We found that a 50µM concentration of Rockout did not disrupt normal cell proliferation and that 100% of 38 hpf-treated embryos survived up to two days in a continuous treatment (Supplementary Fig. 6). Embryos cultured in 50% FASW alone at 26°C, or with the equivalent amount of DMSO, hatched normally (Supplementary Fig. 6).

### Microscopy

Embryos for multiview lightsheet imaging were injected at the 1-cell stage with 2 µg/µl *H2A-mCherry* (nuclei) and *Lyn-EGFP* (membrane) mRNA reporter constructs. We mounted 4-cell stage embryos inside of a 2mm PTFE tube and 3mm glass capillary suspended in 1% low melt agar made in filtered ASW and infused with 500 nm fluorescent beads (F-X050, F-Y050 or F-XC050 microspheres, Estapor Merck). Capillaries were mounted on a SiMView multiview lightsheet microscope^103^ and embryos imaged with a 488 nm (*Lyn-EGFP*) and 594 nm (*H2A-mCherry*) laser at ten-minute intervals from three or four angles with two objectives positioned 180° apart. The z-interval was set to 0.41µm. The resulting six or eight views per sample were registered and fused using the MultiView Reconstruction and BigDataViewer plugins for Fiji^104–107^.

Live imaging of *H2B-mRFPruby* transgenic embryos and *LA-mNG* or *Ph-sqh-mNG* mRNA injected embryos was carried out at 26°C on a Nikon CSU-W1 SoRa Spinning Disc confocal microscope with a 20x objective. Embryos were mounted individually on glass microscope slides so that a 1.5 coverslip with clay-feet spacers could be used to roll the desired region of the germ-disc or grid into the field of imaging. We applied slight pressure to the coverslip to flatten the tissue and obtain clearer views of apical cell face geometries. We then filled the space between the coverslip and slide with FASW and sealed the edges with Vaseline to prevent evaporation. Time-lapse imaging of live embryos was carried out at ten-minute intervals with a z-interval of 0.9µm. Fixed and stained embryos were cleared in 70% glycerol and mounted individually in the same manner as live embryos for confocal imaging on either a Nikon CSU-W1 SoRa Spinning Disc (0.9µm z-interval) or Zeiss LSM 880 (1µm z-interval). Only timelapse datasets taken of embryos that hatched successfully were used for analysis.

### Image analysis and segmentation

Lineage tracing of the fused lightsheet datasets was performed with either semi-automated (Embryo 1; Supplementary Table 1) or manual tracking (Embryos 2 and 3). The nuclear channel of Embryo 1 was first pixel classified and then object tracked in ilastik^108^ (ver. 1.3.3), followed by manual correction in MaMuT^94^ (ver. 1.53), a plugin for Fiji. Embryos 2 and 3 were tracked entirely manually using both the nuclear and membrane channels in MaMuT ^94^ or Mastodon in Fiji. We created the 2D conformal maps of the embryo’s surface from the lightsheet datasets using a version of ImSAnE^109^ customized for *P. hawaiensis*^34^. Nuclear tracks colored according to 16-cell stage lineages were superimposed onto the maps using a custom MATLAB script or manually superimposed in Adobe Illustrator (CC 2023 or 2024). Maximum intensity projections of the lightsheet 2D conformal maps of *Lyn-EGFP* (membrane marker) were segmented in the Fiji plugin TissueAnalyzer^110^ using a watershed algorithm with manual correction.

To visualize and quantitatively describe cell shapes, we first pixel-classified membranes in 10µm maximum intensity projections of Phalloidin-stained or *LA-mNG*-labeled embryos in ilastik^108^ (ver. 1.4.0) to generate a more consistent cell boundary signal. Especially in the later grid stages (approximately 62 hpf onwards), the embryo must be carefully laid flat against the slide and slightly compressed with the coverslip. For samples in which flattening out all wrinkles or depressions in the tissue was not possible, the probability maps containing apical cell faces for each tissue region were cut out and composited together using the Image Calculator in Fiji. We then fed the probability maps exported from ilastik^108^ (ver. 1.3.3) into Cellpose^111^ (ver. 3.0) and obtained cell ROIs using a custom-trained model with some manual correction of contiguous masks. Automatic size calibration was performed and non-contiguous or incomplete cell masks around the edges were removed. All subsequent quantitative descriptions of cell shape and tissue organization were carried out on these ROIs using custom Python scripts available at https://github.com/boyonpointe/parhyale_morpho/tree/main with commit ID a19a88d. All other measurements, including intensities of cytoskeletal components, were made on 10µm maximum intensity projections in Fiji (ver. 2.1.0) using the ROI section and associated measurement tools.

For cell shape and vertex IL measurements, each 10 μm maximum intensity projection was cropped to 207 μm wide × 104 μm high. The crop box was centered on the germ disc parallel to the dorso-ventral axis and centered on the midpoint of the IL (a clear straight or slightly bent line) in the anterior-posterior axis. Only cells contained fully within the crop box were counted.

For IL actin and myosin 2 measurements, in un-cropped 10 μm maximum intensity projections, the lateral extent of the IL region of interest was determined by drawing a line from the ventral midline of the germ disc towards the left and right sides of the germ disc. The region was considered terminated when it reached cells whose sides were bent more than approximately 15° from the leading edge of the region. The mean IL length for phalloidin-stained samples (F-actin) was 146 μm (n = 38) and the mean IL length for *Ph-sqh*-labeled samples (myosin 2) was 151 μm (n = 14).

To compare cell shape distributions in the posterior of the germ disc, we used a bootstrap approach. For each pair of distributions, we generated 100,000 surrogate pairs of random resampling and calculated the difference in their means. This gave us a null distribution of differences, against which we assessed how often the observed empirical difference would occur.

The p-value corresponds to that probability. For stop motion in Movie 3, images were printed on light-permeable paper and animated over a lightbox using a Sony A7S camera controlled by the stop-motion animation software DragonFrame (v4).

### Cell shape descriptor

To measure cell shape, we used a custom descriptor in which the area of a cell face is divided by the area of the circle whose diameter is equal to the distance between the farthest two points on the cell’s perimeter. With this descriptor, a perfect square has a value of 0.64, and hexagons and higher order polygons have values of approximately ⩾0.75. We found this measurement to more usefully describe the emergence of square-cell packing than counting the number of cell sides, because within this dynamic tissue, 4-way vertices frequently dissolve into small edges at square corners, and cell divisions can introduce temporary kinks into neighboring cell edges.

### Cellular Potts Model Implementation

Our model to test the effect of differential adhesion between different cell lineages was formulated in the Cellular Potts Model (CPM)^41,67,112^. We implemented our model using Compucell3D^43^. In CPM, the cells are modeled as a spatially contiguous collection of pixels. The pixels that are at the boundary of cells have an energy cost associated with interacting with pixels belonging to the neighboring cells. The energy cost between a pair of pixels (a,b) depends on the cell type (σ(a), σ(b)) of both the cells to which the interacting pair of pixels belong. The area of (A) of a cell is defined as the number of pixels the cell encompasses while the perimeter (L) is the number of pixels at the boundary of the cell.

The following equation shows the expression of energy of the system for given pixel configuration:

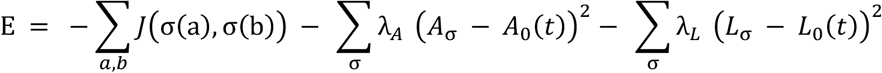

The second and third terms on the right hand side of the equation represent the energy cost for cell area and cell perimeter to deviate away from the steady state values *A*_0_ and *L*_0_ respectively. To implement growth, *A*_0_ is made a function of time such that the cell grows to reach *A*_0_ and when it meets the size to undergo division, it divides. The steady state value for the perimeter is also a function of time and set to ensure that steady state shape is circular i.e. 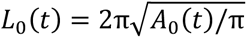. The symmetric matrix *J* has its number of rows and columns equal to the number of cell types in the system. Each entry *J_i,j_* in the matrix specifies the interaction energy between pixels belonging to cell type i and j. In our simulation there are six different cell types corresponding to six different lineages and an additional one that corresponds to the medium.

The segmented cell masks of tissue imaged at 26 hpf is used to generate the initial configuration of the pixels in the simulation. Each pixel belongs to one among the 6 different cell types or to the medium. The cells increase in size by 100 pixels before they divide. The amount of growth cells undergoes before they divide is chosen to reflect the empirically observed increase in tissue size and cell numbers seen at 46 hpf. The cells divide symmetrically along the axes orthogonal to the major axis (long axis). Throughout the simulation, we track the number of pixels in contact with pixels from cells of different lineages.

### Ablation experiments

Ablation experiments were carried out on a Zeiss 980 NLO 2-photon microscope with a pulsed Spectra-Physics InSight X3+ laser set to 927 μm wavelength and an ablation ROI with a 1µm width that was passed over five times. *LA-mNG* embryos injected at the 1-cell stage were incubated to the desired stage and mounted individually on slides with a Vaseline-sealed coverslip. Ablated embryos were imaged with a LD LCI Plan-Apochromat 25x/0.8 objective for two frames prior to the ablation followed by every second for two minutes immediately afterwards. Some embryos were further imaged on the same microscope at intervals of one minute to follow the effects of the ablation. Embryos used for further analysis were then recovered from the slide and transferred to a 35mm petri dish with fresh FASW with 1:100 Penicillin-Streptomycin (10,000 U/mL penicillin, 10,000 µg/ml streptomycin; Gibco 15140-122) and 1:200 Amphotericin-B (250 µg/ml, Gibco 15290-026) for incubation at 26 °C until fixation. Recoil length was calculated in Fiji by subtracting the distance between cell vertices on either side of the ablation ROI in the first frame immediately post ablation, from the distance between the same two vertices before the incision was made.

### Figure presentation

Line tracing of membranes shown in figures was performed using the Image Trace function (Silhouettes or Line Art) on exported Cellpose^111^ ROIs or on hand segmented micrographs (Supplementary Fig. 3) in Adobe Illustrator 2024 or 2025. To more clearly display the relationship between Engrailed expression and grid geometry in Figure 1, we used ilastik^108^ (ver.1.3.3) to pixel-classify images of anti-Engrailed staining and exported the resulting probability maps. In Movie 4, TissueAnalyzer^113^ was used to track and visualize cell membranes of the midline founder population in a conformally mapped time-series that was manually aligned. All figures were assembled with Adobe Illustrator.

## Supporting information

Movies 1 through 4

## Acknowledgements

The authors thank William Lemon and Philipp Keller (HHMI Janelia Research Campus) for providing access and training on their SimView lightsheet microscope, Sebastian Streichan and Dillon Cislo (University of California, Santa Barbara) for help with ImSAnE and applying MaMUT tracks to ImSAnE maps, John Rallis and Myrto Ziogas (IMBB-FORTH) for help with plasmid preparations, as well as members of the Extavour and Pavlopoulos laboratories for discussion.

## Author Contributions

BS conceived of the study, designed experiments, performed all wet lab experiments, performed data analysis and interpretation, and wrote the first draft of the manuscript. LB designed, performed embryo collection and injection experiments for, and executed capture of lightsheet datasets, and reviewed the manuscript. CK designed and performed all computational modelling experiments, performed data analysis, and reviewed the manuscript. ES and AP conceived of the study, designed and created all fluorescent *P. hawaiensis* constructs, performed embryo collection and injection experiments for capture of lightsheet datasets, and reviewed the manuscript. CGE conceived of the study, performed data interpretation, obtained funding for the study, supervised its execution, and edited the manuscript.

## Data Availability

Lightsheet datasets described in Supplementary Table 1. MaMuT lineage tracks for these datasets, as well as scripts for cellular potts model simulations and image analyses, are available at https://github.com/boyonpointe/parhyale_morpho/tree/main with commit ID a19a88d.

## Statement on Use of Machine Learning, Artificial Intelligence and Large Language Models

ChatGPT 4.0 and 5.0 (open AI) were used to modify R code for altering plot aesthetics. ilastik^108^ and Cellpose^111^, used for image segmentation in this study, are machine learning-based programs. ML/AI was not used to generate any content, write any text, or perform any other data analysis.

## Movie Legends <u>Download Link</u>

All movies show Embryo 1 (see Supplementary Table 1).

**Movie 1**. **The first 2.5 days of development of a membrane-labeled *P. hawaiensis* embryo**. Timelapse animation of the *Lyn-EGFP* (membrane) channel of a reconstructed multiview lightsheet dataset beginning at the 8-cell stage and ending during germ band grid elaboration. Each frame is a maximum intensity projection through half of the embryo and was taken ten minutes apart. 319 total frames are animated at a frame-rate of nine frames per second. Sequential frames from this animation were selected for and sequentially displayed in Figure 1 (first frame is Fig. 1b).

**Movie 2**. **The first 2.5 days of development of a nuclear-labeled *P. hawaiensis* embryo**. Timelapse animation of the *H2A-mCherry* (nuclei) channel of the same embryo shown in Movie 1. Each frame is a maximum intensity projection through half of the embryo and was taken ten minutes apart. 319 total frames are animated at a frame-rate of nine frames per second.

**Movie 3**. **Dynamics of the 16-cell stage lineage domains visualized by stop-motion animation**. The left side of this animation shows a still from Movie 1 with the *Lyn-EGPF* marked 16-cell stage ectodermal blastomeres false colored (Era = magenta, Ela = pink, Erp = yellow, Elp = purple, Erp = cyan, Elp = orange). The right side shows 2D maps of the same dataset in which the MaMuT nuclear tracks have been superimposed on the membrane channel to show the lineage of each cell. The rectangular cut-out corresponds to the cropped regions of each map shown in Fig. 2c, Fig. 6a, Supplementary Fig. 1 and Supplementary Fig. 8a. The maps shown here begin at 42 hpf and end at 55.67 hpf and are animated at a frame rate of nine frames per second, displaying 1.5 hours of developmental time every second. Given that the mapping of the 3D surface to 2D was not constrained and the germ disc migrates across the yolk, frames were manually aligned to produce animated motion and keep the emerging grid fixed within the cut-out region. The first sweep of the pointing finger across the map (00:07) indicates the IL (Fig. 2) at the anterior-posterior compartment boundary. The second sweep at the end (00:10) indicates the emerging midline (Fig. 4), which separates the Erp and Elp lineages by one cell beginning just posterior to the IL (P1 in Fig. 4b).

**Movie 4. Intercalation of midline cells to form a column of square cells.** The midline of the germ band grid forms as a single column of cells via intercalation. Cells aligned into the midline and within the intercalation zone at 66±2 hpf were tracked backwards to the onset of midline formation.

**Supplementary Figure 1.**
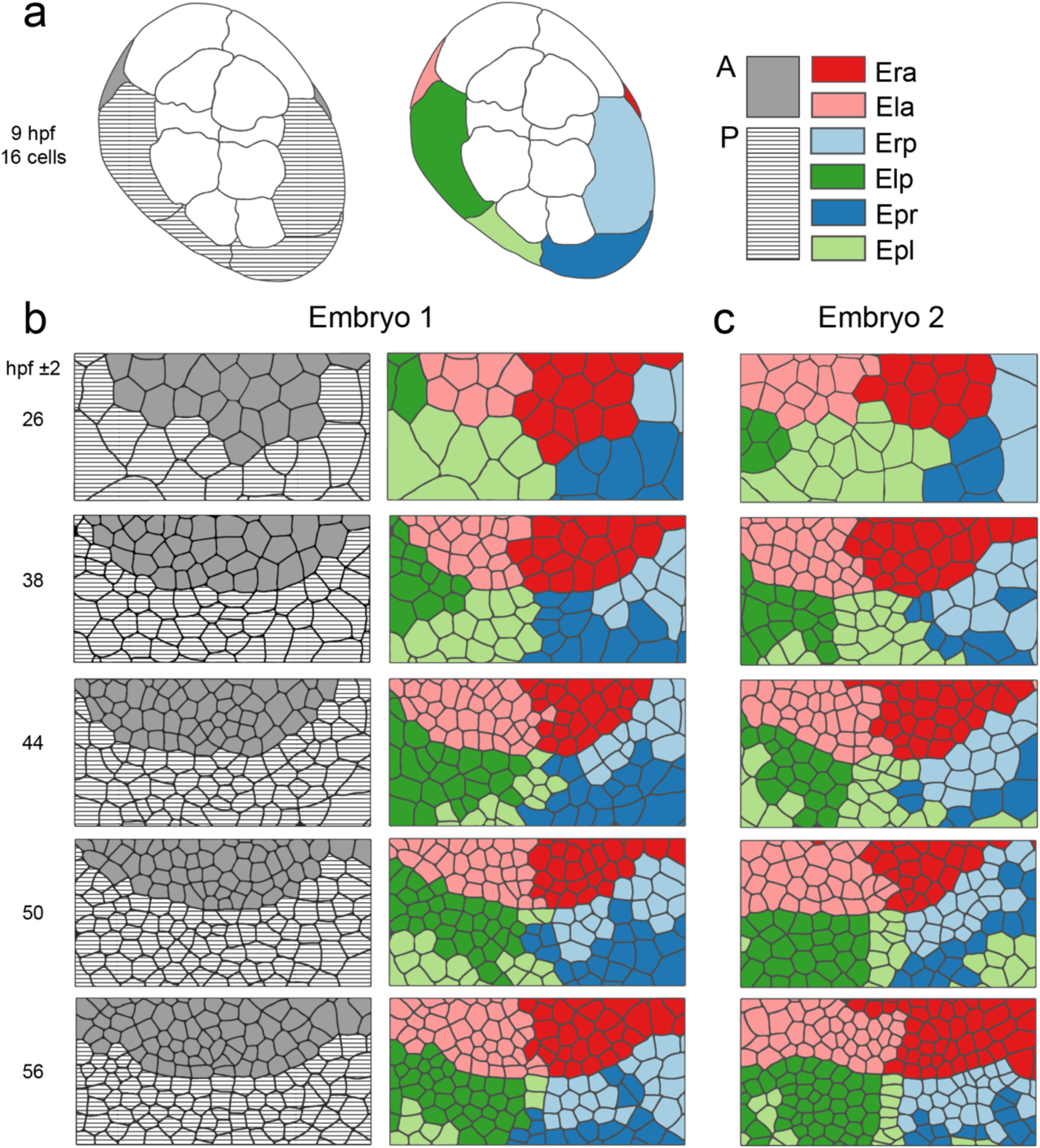
16-cell ectodermal lineage domains undergo extensive rearrangement. Stills from long-term multiview lightsheet timelapses shown either as tracings of maximum intensity projections (a) or 2D conformal maps (b-c) in which lineage was determined by tracking each cell back to its 16-cell stage ancestor. **a** 16-cell ectoderm precursor blastomere identities for Embryo 1 (Supplementary Table 1) colored with reference to anterior (grey)-posterior (striped) compartments or 16-cell stage identity (colors). **b-c** Lineage domain comparison between Embryo 1 and Embryo 2 (Supplementary Table 1) showing that in both embryos, anterior lineages (Era, Ela) do not mix with posterior lineages (Erp, Elp Epr, Epl). In the posterior compartment, the Erp and Elp domains begin on opposite (left and right) sides of the germ disc at 26±2 hpf and migrate closer to one another until they are separated by only a single column of Epr or Epl cells by 56±2 hpf, indicating the presence of extensive cell rearrangement.

**Supplementary Figure 2.**
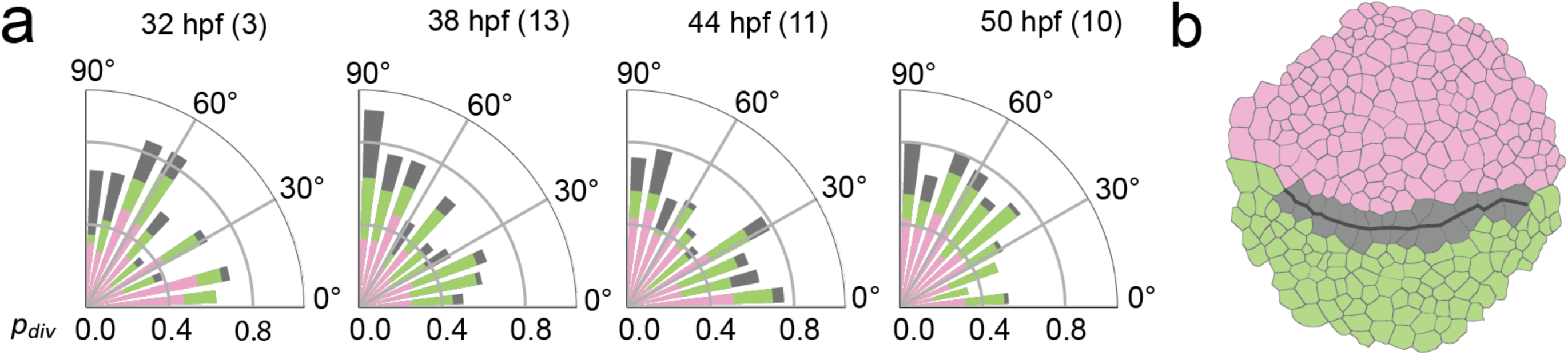
Cell proliferation occurs throughout the germ disc from 32-30 hpf, with IL-adjacent divisions oriented preferentially perpendicular to the IL. **a** Cell division angles measured across the germ disc in 32-50 hpf embryos (sample size in parentheses). Angles are plotted with reference to the orientation of the IL (0°) and colored according to the division region (anterior = pink, IL = grey, posterior = green). **b** Segmented germ disc indicating locations of defined tissue regions. IL cell divisions are those that occur in cells directly adjacent to the IL on the anterior or posterior side (grey).

**Supplementary Figure 3.**
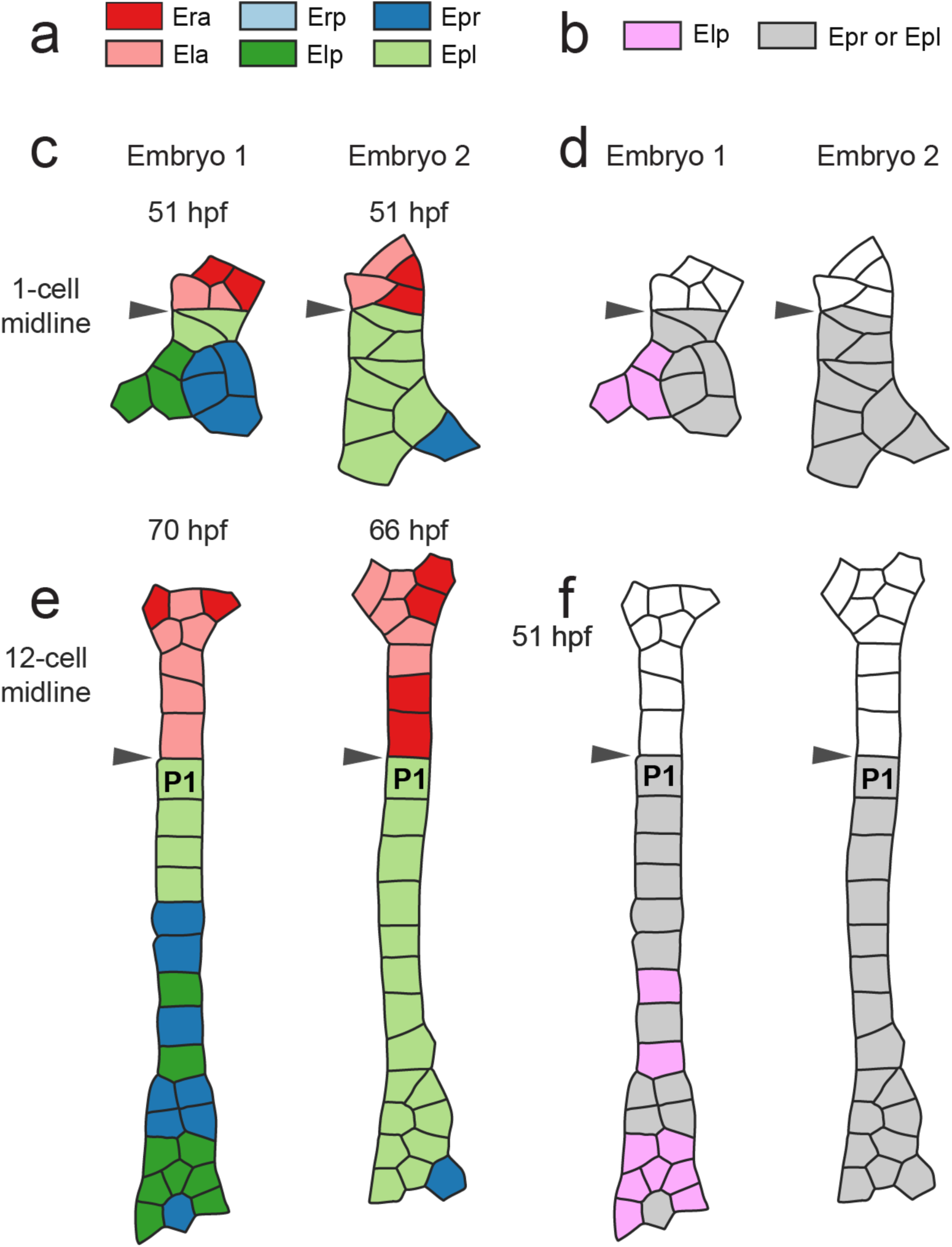
Multiple 16-cell stage ectodermal lineages contribute to the midline, including non-Ep clones in the posterior. **a** Legend for 16-cell stage ectodermal precursor blastomeres. Midline descendants shown in c and e. **b** Legend for 16-cell stage Ep and non-Ep ectodermal precursor blastomeres. **c, e** The midline is composed of descendants of multiple 16-cell blastomeres both anterior and posterior to the anterior-posterior compartment boundary, which we call the IL at earlier stages. Midline cells in 12-cell midline embryos were tracked back to the 16-cell stage in the lightsheet datasets. At the 1-cell midline timepoint, only cells whose descendants make up the midline or intercalation zones shown in the 12-cell midline timepoints, are shown. **d, f** The posterior midline is not exclusively made up of descendants of Ep. Arrowheads indicate the location of the compartment boundary. In c through f, Embryos 1 and 2 correspond to the same datasets shown in Supplementary Figure 1 and described in Supplementary Table 1.

**Supplementary Figure 4.**
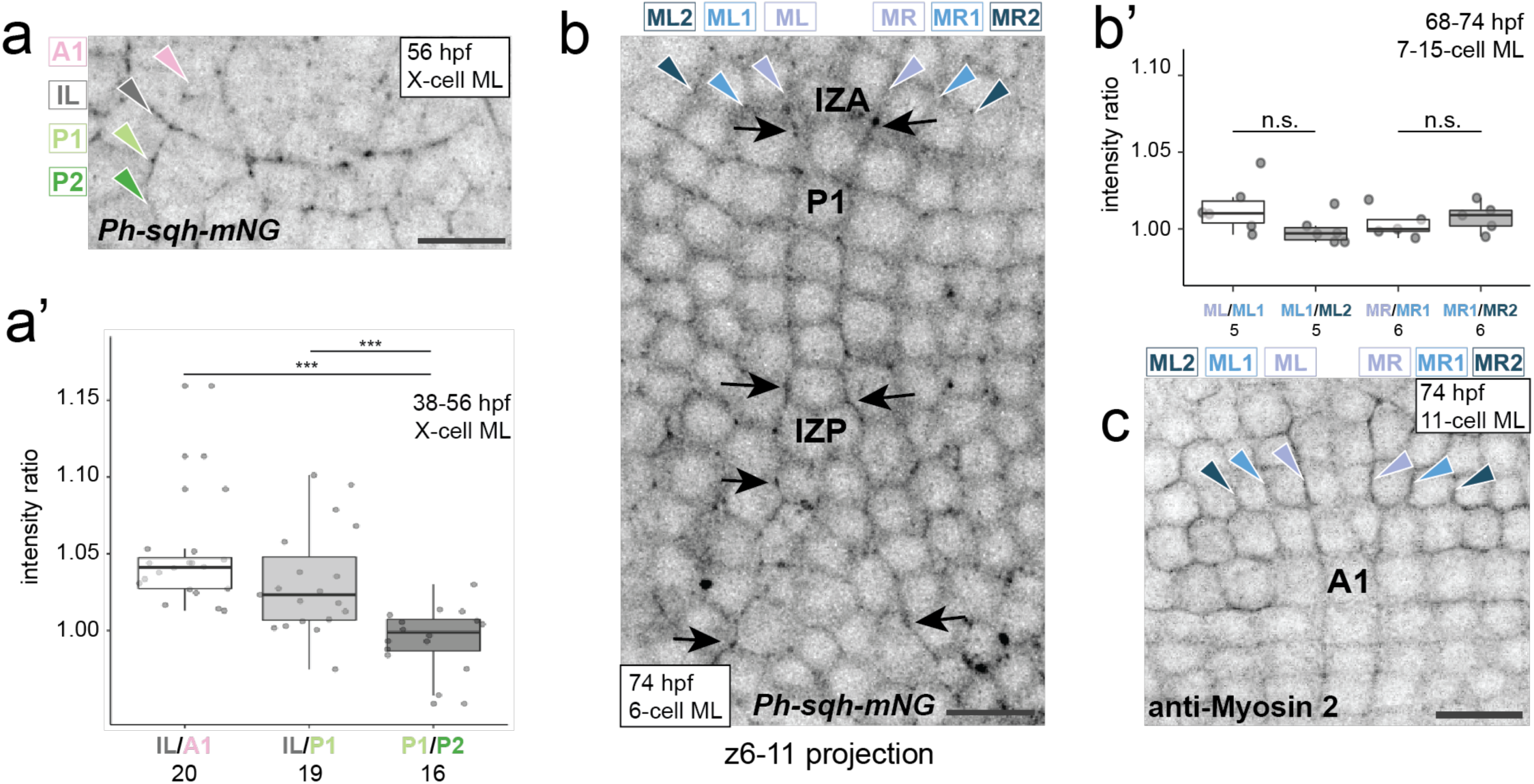
Myosin 2 accumulates at the IL and midline cables. **a** 10µm maximum intensity projection of a 56±2 hpf embryo injected with *Ph-sqh-mNG* mRNA to label the myosin 2I regulatory light chain. **a’** Ratios of mean intensity of *Ph-sqh-mNG* signal measured from 38-56 hpf embryos in 10µm maximum intensity projections like that shown in a. The regions where *Ph-sqh* signal was quantified (A1, IL, P1, P2) are shown in a; each begins at the arrowhead and extends across contiguous cell sides to the rightmost edge of the region shown. **b** Maximum intensity projection through 5µm of a 74 hpf embryo at the 6-cell midline stage. Arrows indicate regions of myosin- 2 accumulation along the MR and ML cables (arrowheads), and at the anterior and posterior intercalation zones (IZ). **b’** Ratios of mean intensity of *Ph-sqh-mNG* signal measured from 5µm maximum intensity projections labeled with *Ph-sqh-mNG* expression. The regions quantified (MR, MR1, MR2, ML, ML1, ML2) are shown in b; each begins at the arrowhead and extends down contiguous cell sides to the bottom of the region shown. p = 0.05 for ML/ML1 compared to ML1/ML2; p = 0.20 for MR/MR1 compared to MR1/MR2. **c** 5µm maximum intensity projections of a 74 hpf embryo with 11 cells at the midline, labeled with an anti-Myosin 2 antibody. Only the anterior portion of the midline is shown here. Scale bars = 25µm and anterior is up in all micrographs. For all statistical comparisons * p-value < 0.05, ** p-value < 0.01, *** p-value < 0.001, and n.s. = not significant (Student’s t-test for a; Mann Whitney U-test for b; all tests one-tailed). Sample sizes in a’ and b’ (numbers of embryos) indicated by numbers under category labels.

**Supplementary Figure 5.**
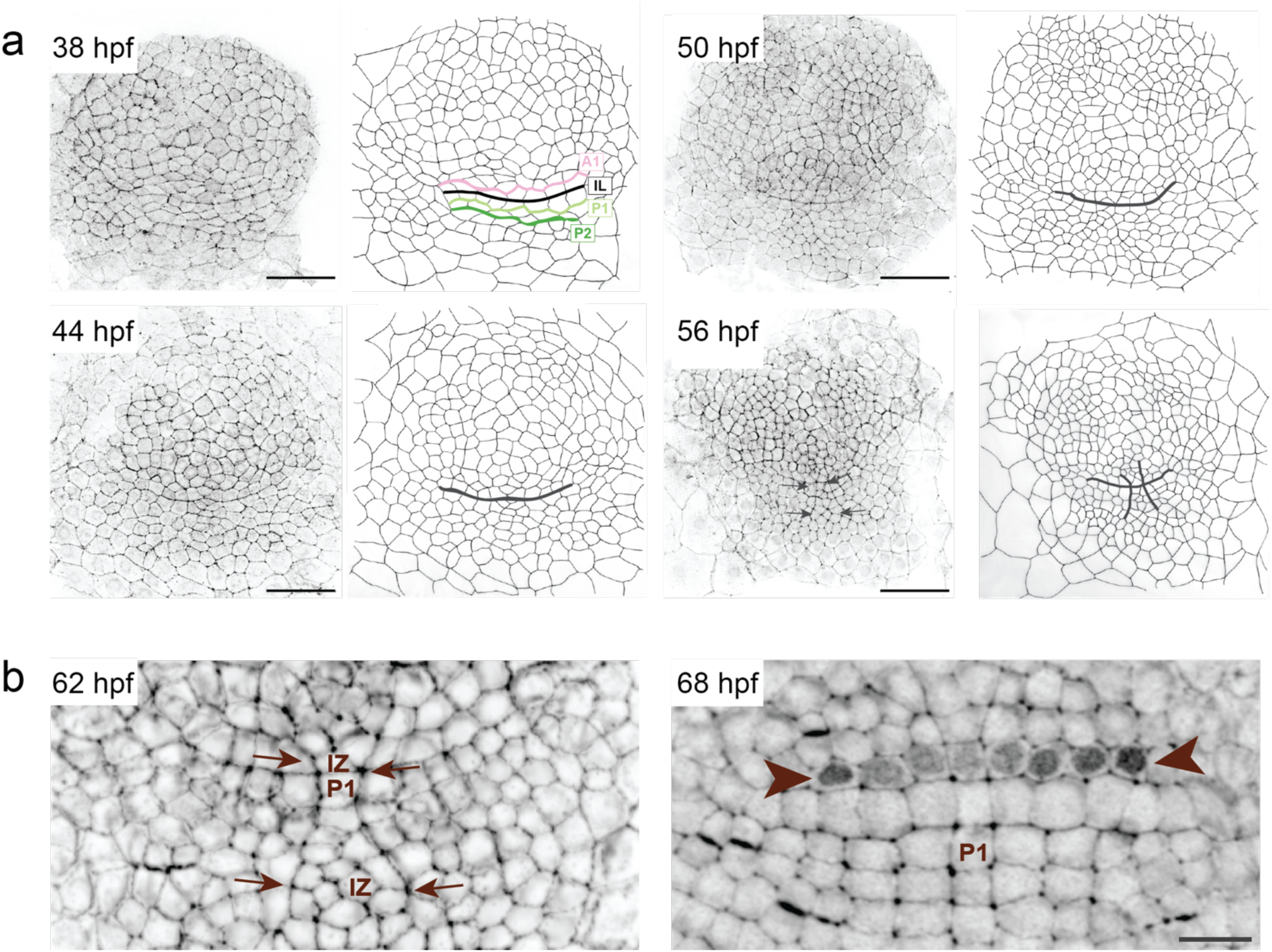
IL and midline cables in the context of the whole germ disc. **a** 10µm maximum intensity projections of 38-56±2 hpf embryos stained with phalloidin (left) and with membranes traced (right). IL (all stages shown) and midline cables (56 hpf only) demarcated with thick lines. Anterior is up; scale bar = 100µm. **b** 10µm maximum intensity projections of the regions around the IL and midline cables in 62-68±2 hpf embryos. Arrows indicate the midline cables flanking the intercalation zone (IZ). Arrowheads indicate Engrailed expression in the prospective mandibular segment (Mn). Anterior is up; scale bar = 20µm.

**Supplementary Figure 6.**
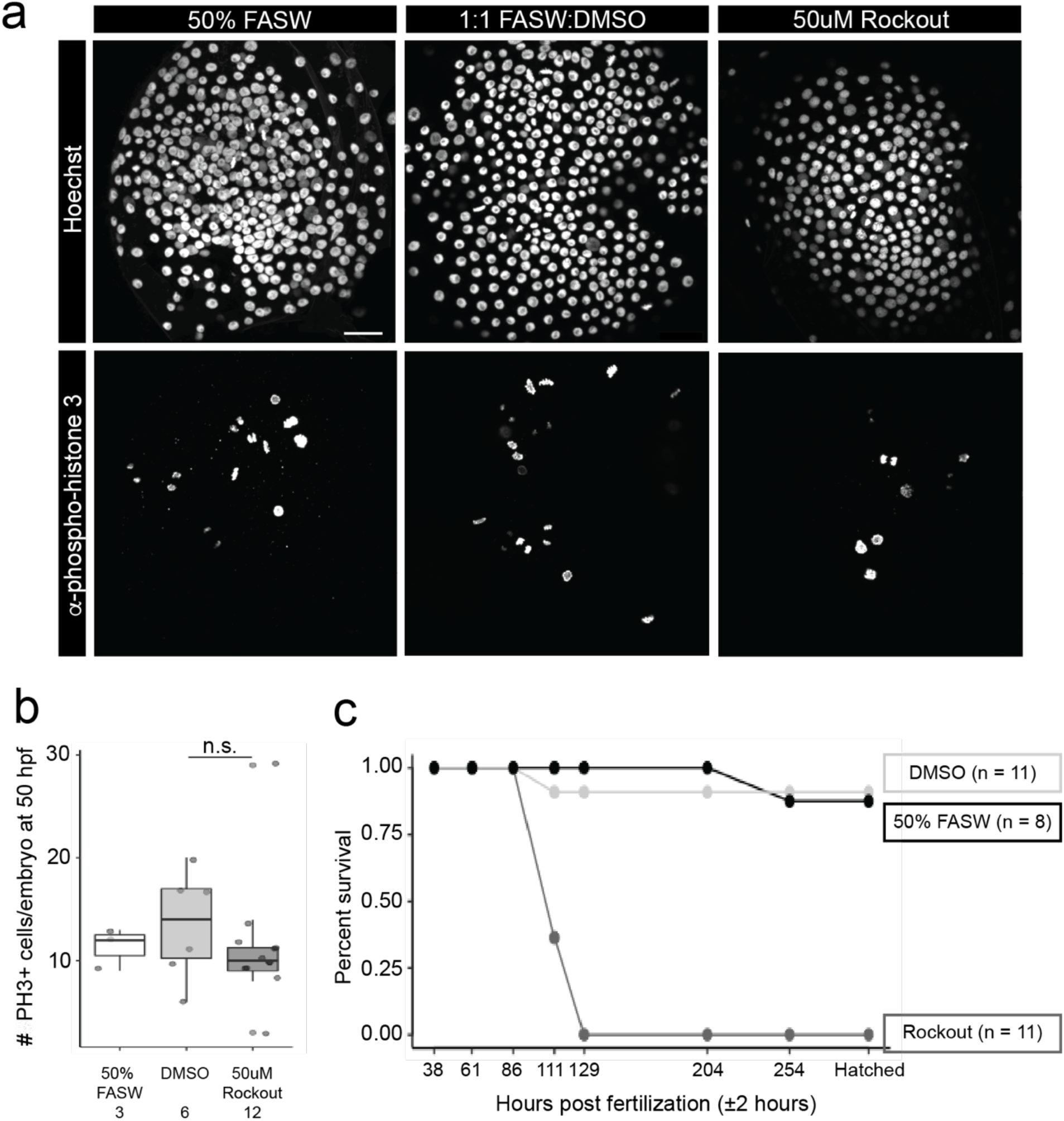
12-hour Rockout treatment does not impact cell proliferation. **a** 38±2 hpf embryos treated with either 50% FASW, 1:1 FASW:DMSO, or 50µM Rockout in 50% FASW were fixed 12 hours later (at 50±2 hpf), stained with Hoechst 33342 (top row) and anti-phospho-histone H3 (bottom row). Scale bar = 100µm and applies to all panels. **b** Total number of phospho-histone H3-positive cells for each treatment shown in a. n.s. = not significant (p-value = 0.303, two-tailed Mann-Whitney U Test). **c** Continuous treatment hatching assay. 100% of embryos incubated in 50µM Rockout continuously beginning at 38±2 hpf remained alive after 48 hours of treatment (86±2 hpf), and all died by 91 hours of treatment (129±2 hpf) (n = 11 embryos). By contrast, 100% of embryos in the 50% FASW (n = 8) and 1:1 FASW:DMSO (n = 11) treatments remained alive until 204±2 hpf. 87.5% hatched in 50% FASW and 90.1% of embryos hatched in 1:1 FASW:DMSO. Sample sizes in b indicated by numbers underneath category labels.

**Supplementary Figure 7.**
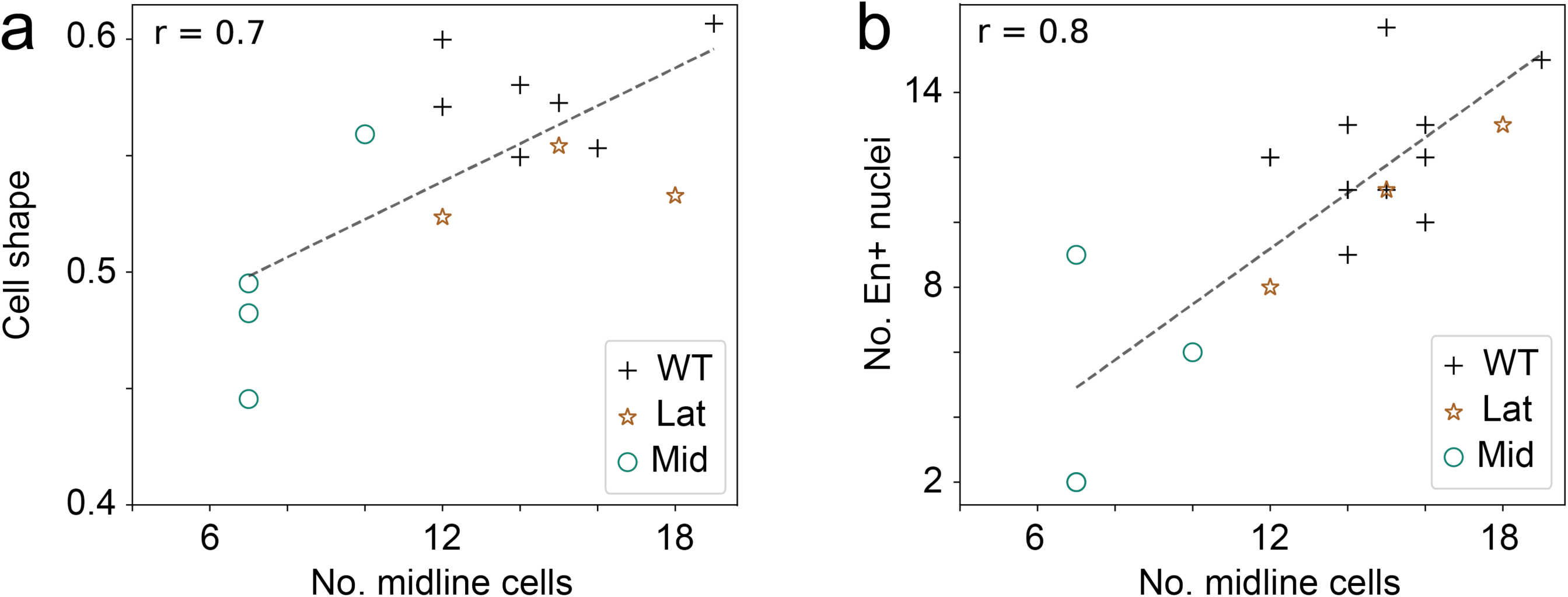
**a** The bottom tenth percentile of the cell shape value (Fig. 1l) distribution of the whole tissue (i.e. the most anisotropic/least round cells) correlates positively with the total number of midline cells for that sample, in both wild type and ablated embryos. For wild type embryos, the roundness value of the bottom tenth percentile of cells is close to 0.6, implying that the shape of most non-round cells in wild type tissues is a square. However, upon midline ablation, the roundness value for this bottom tenth percentile is well below 0.6, implying that cells that would have been square in the wild type adopt more asymmetric shapes. The most dramatic impact on cell shape is experienced by the cells in the midline, which are approximately 10% of the cells in the tissue at this stage (data not shown). **b** The total number of Engrailed-positive (En+) nuclei correlates positively with the total number of midline cells.

**Supplementary Figure 8.**
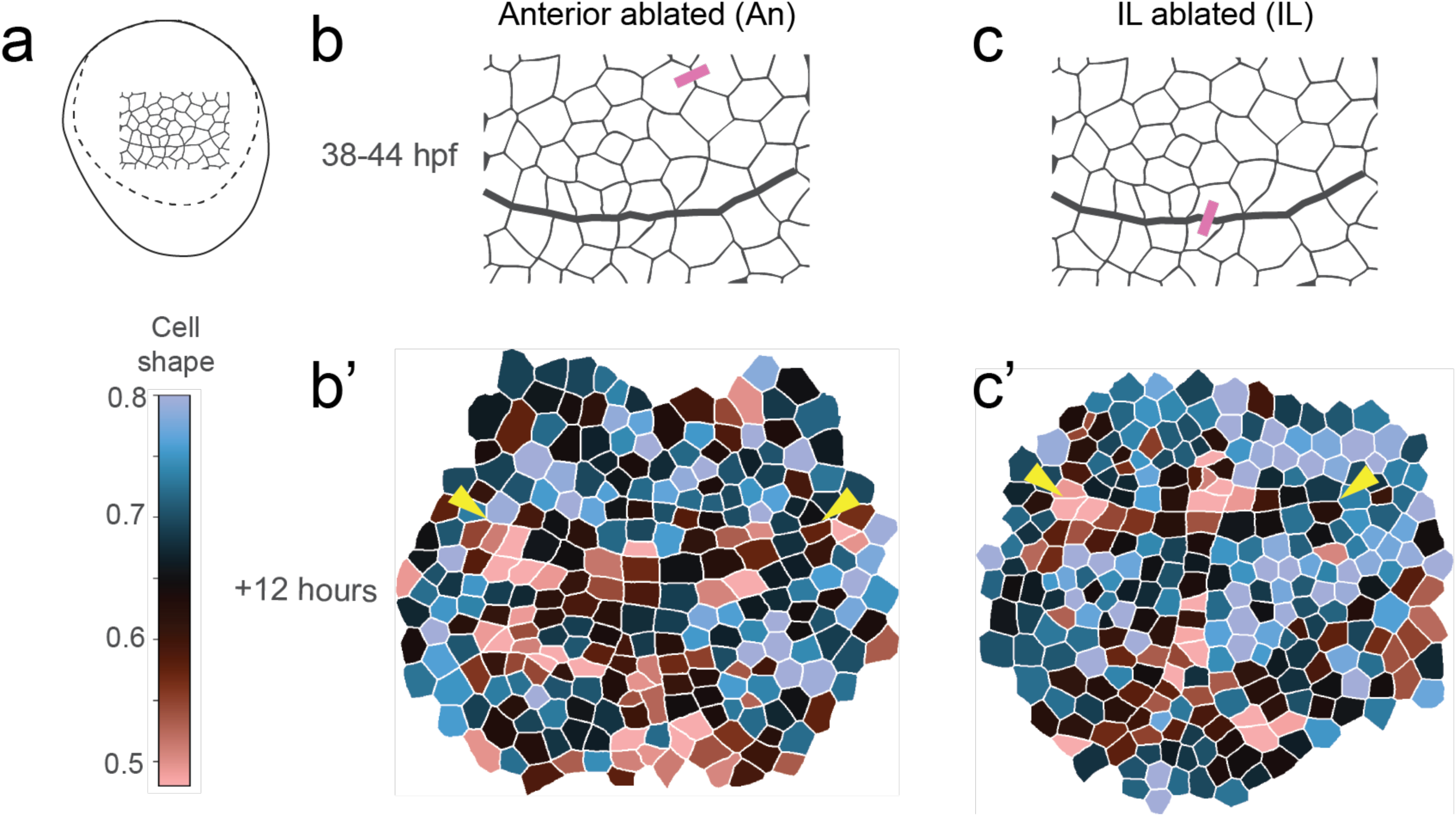
Ablated IL embryos maintain a straight line at the IL and form a midline. **a** Schematic showing the location of the region delineated in b and c. **b** Ablation of a cell bond four cells anterior to the IL (n = 2). Ablation sites indicated by the pink rectangles. **b’** Cell shape descriptor values mapped on to +12 hour ablation tissue segmentations. **c** Ablation of a cell bond at the IL (n = 5). Ablation site indicated by the pink rectangle. **c’** Cell shape descriptor values mapped as described in b’. Arrowheads: IL.

**Supplementary Table 1.** Lightsheet datasets used for analyses. h: hours. min: minutes.

| **Lightsheet Sample** | **Total imaging time (h)** | **T-interval (min)** | **16-cell lineages tracking** | **Midline tracking** | **Posterior cell numbers** | **Cell division orientation** | **Tissue morphology seeds for simulation** |
| --- | --- | --- | --- | --- | --- | --- | --- |
| Embryo 1 | 61.00 | 10 | Yes | Yes | Yes | Yes | Yes |
| Embryo 2 | 61.00 | 10 | Yes | Yes | Yes | Yes | No |
| Embryo 3 | 66.33 | 10 | No | Yes | Yes | Yes | No |
| **Figure** |  |  | **Fig. 2, Supp. Fig. 1** | **Fig. 4, Supp. Fig. 2** | **Fig. 5** | **Supp. Fig. 3** | **Fig. 3** |

**Supplementary Table 2.** Mitotic activity in the germ rudiment at the time of intercalation line (IL) formation. “Live” samples correspond to lightsheet sample embryos (Supplementary Table 1). Data from samples WT1, WT2 and WT3 were used to generate rose plots in Supplementary Figure 2a.

|  |  |  |  |  | Live: 6hr time window |  |  |  | Fixed |  |  |  |  |  |  |  |
| --- | --- | --- | --- | --- | --- | --- | --- | --- | --- | --- | --- | --- | --- | --- | --- | --- |
|  |  |  |  |  | Cell divisions in germ disc |  |  |  | Cell divisions in germ disc |  |  |  | Total cells in germ disc |  |  |  |
| hpf | N (live) | N (fixed) | N (total) |  | Total | A | IL | P | Total | A | IL | P | Total | A | IL | P |
| **32** | 3 | 0 | 3 | Mean | 54.33 | 31.33 | 9.67 | 13.33 | 4.67 | 3.33 | 1.33 | 0.00 | 154.00 | 84.00 | 16.67 | 53.33 |
|  |  |  |  | SD | 17.16 | 14.05 | 4.04 | 1.53 | 1.53 | 2.31 | 1.53 | 0.00 | 18.19 | 14.93 | 1.15 | 13.65 |
|  |  |  |  | SE | 9.91 | 8.11 | 2.33 | 0.88 | 0.88 | 1.33 | 0.88 | 0.00 | 10.50 | 8.62 | 0.67 | 7.88 |
| **38** | 3 | 10 | 13 | Mean | 69.33 | 30.33 | 13.00 | 26.00 | 5.42 | 2.67 | 1.00 | 1.92 | 238.92 | 155.33 | 20.62 | 64.31 |
|  |  |  |  | SD | 8.50 | 5.13 | 4.58 | 1.73 | 2.11 | 1.61 | 1.00 | 1.04 | 23.60 | 20.72 | 2.10 | 7.97 |
|  |  |  |  | SE | 2.36 | 1.42 | 1.27 | 0.48 | 0.58 | 0.45 | 0.28 | 0.29 | 6.55 | 5.75 | 0.58 | 2.21 |
|  |  |  |  | A vs P |  |  |  |  | 0.10565 |  |  |  | **<0.00001** |  |  |  |
| **44** | 2 | 10 | 12 | Mean | 108.50 | 66.00 | 16.00 | 26.50 | 5.45 | 2.91 | 0.75 | 1.75 | 311.91 | 183.64 | 24.33 | 103.83 |
|  |  |  |  | SD | 21.92 | 7.07 | 8.49 | 6.36 | 2.62 | 2.07 | 0.75 | 1.36 | 31.23 | 19.30 | 2.57 | 25.56 |
|  |  |  |  | SE | 6.33 | 2.04 | 2.45 | 1.84 | 0.76 | 0.60 | 0.22 | 0.39 | 9.02 | 5.57 | 0.74 | 7.38 |
|  |  |  |  | A vs P |  |  |  |  | 0.09342 |  |  |  | **0.00006** |  |  |  |
| **50** | 2 | 9 | 11 | Mean | 141.00 | 79.00 | 15.50 | 46.50 | 5.33 | 3.33 | 0.36 | 1.67 | 356.22 | 211.11 | 24.82 | 120.11 |
|  |  |  |  | SD | 22.63 | 18.38 | 0.71 | 4.95 | 2.45 | 1.87 | 0.50 | 1.12 | 56.18 | 36.46 | 2.48 | 24.14 |
|  |  |  |  | SE | 6.82 | 5.54 | 0.21 | 1.49 | 0.74 | 0.56 | 0.15 | 0.34 | 16.94 | 10.99 | 0.75 | 7.28 |
|  |  |  |  | A vs P |  |  |  |  | **0.03216** |  |  |  | **0.00029** |  |  |  |

**Supplementary Table 3.** En expression in cells of the prospective mandibular segment in wild type (WT) embryos, following ablation of cell bonds ral to the midline (Lat), or following ablation of cell bonds at the midline, as depicted in Fig. 6b, c. These data were used to generate Fig. 6e.

|  |  | Row cell identity |  |  |  |  |  |  |  |  |  |  |  |  |  |  |  |  |
| --- | --- | --- | --- | --- | --- | --- | --- | --- | --- | --- | --- | --- | --- | --- | --- | --- | --- | --- |
| Experiment | Sample # | L7 | L6 | L5 | L5 | L3 | L2 | L1 | ML | R1 | R2 | R3 | R4 | R5 | R6 | R7 | Total # En+ | Sample Mean |
| Wild Type | 1 | 0.00 | 0.00 | 0.00 | 1.00 | 1.00 | 1.00 | 1.00 | 1.00 | 1.00 | 1.00 | 1.00 | 1.00 | 0.00 | 0.00 | 0.00 | 9 | 0.60 |
|  | 2 | 0.00 | 0.00 | 1.00 | 1.00 | 1.00 | 1.00 | 1.00 | 1.00 | 1.00 | 1.00 | 1.00 | 1.00 | 1.00 | 0.00 | 0.00 | 11 | 0.73 |
|  | 3 | 0.00 | 0.00 | 1.00 | 1.00 | 1.00 | 1.00 | 1.00 | 1.00 | 1.00 | 1.00 | 1.00 | 1.00 | 1.00 | 0.00 | 0.00 | 11 | 0.73 |
|  | 4 | 0.00 | 0.00 | 0.00 | 1.00 | 1.00 | 1.00 | 1.00 | 1.00 | 1.00 | 1.00 | 1.00 | 1.00 | 0.00 | 0.00 | 0.00 | 9 | 0.60 |
|  | 5 | 0.00 | 0.00 | 1.00 | 1.00 | 1.00 | 1.00 | 1.00 | 1.00 | 1.00 | 1.00 | 1.00 | 1.00 | 1.00 | 0.00 | 0.00 | 11 | 0.73 |
|  | 6 | 0.00 | 0.00 | 0.00 | 1.00 | 1.00 | 1.00 | 1.00 | 1.00 | 1.00 | 1.00 | 1.00 | 1.00 | 1.00 | 0.00 | 0.00 | 10 | 0.67 |
|  | 7 | 0.00 | 0.00 | 0.00 | 1.00 | 1.00 | 1.00 | 1.00 | 1.00 | 1.00 | 1.00 | 1.00 | 1.00 | 0.00 | 0.00 | 0.00 | 9 | 0.60 |
|  | 8 | 0.00 | 0.00 | 0.00 | 0.00 | 1.00 | 1.00 | 1.00 | 1.00 | 1.00 | 1.00 | 1.00 | 0.00 | 0.00 | 0.00 | 0.00 | 7 | 0.47 |
|  | 9 | 0.00 | 0.00 | 0.00 | 1.00 | 1.00 | 1.00 | 1.00 | 1.00 | 1.00 | 1.00 | 1.00 | 1.00 | 0.00 | 0.00 | 0.00 | 9 | 0.60 |
|  | 10 | 1.00 | 1.00 | 1.00 | 1.00 | 1.00 | 1.00 | 1.00 | 1.00 | 1.00 | 1.00 | 1.00 | 1.00 | 1.00 | 0.00 | 0.00 | 13 | 0.87 |
|  | 11 | 0.00 | 0.00 | 0.00 | 1.00 | 1.00 | 1.00 | 1.00 | 1.00 | 1.00 | 1.00 | 1.00 | 1.00 | 0.00 | 0.00 | 0.00 | 9 | 0.60 |
|  | 12 | 0.00 | 0.00 | 0.00 | 1.00 | 1.00 | 1.00 | 1.00 | 1.00 | 1.00 | 1.00 | 1.00 | 0.00 | 0.00 | 0.00 | 0.00 | 8 | 0.53 |
|  | Cell Mean | 0.08 | 0.08 | 0.33 | 0.92 | 1.00 | 1.00 | 1.00 | 1.00 | 1.00 | 1.00 | 1.00 | 0.83 | 0.42 | 0.00 | 0.00 | 9.67 |  |
| Lateral Ablation | 1 | 0.00 | 0.00 | 0.00 | 1.00 | 1.00 | 1.00 | 1.00 | 1.00 | 1.00 | 1.00 | 1.00 | 1.00 | 0.00 | 0.00 | 0.00 | 9 | 0.60 |
|  | 2 | 0.00 | 0.00 | 0.00 | 0.00 | 1.00 | 1.00 | 1.00 | 1.00 | 1.00 | 1.00 | 1.00 | 1.00 | 0.00 | 0.00 | 0.00 | 8 | 0.53 |
|  | 3 | 0.00 | 0.00 | 0.00 | 0.00 | 0.00 | 0.00 | 1.00 | 1.00 | 1.00 | 1.00 | 1.00 | 0.00 | 0.00 | 0.00 | 0.00 | 5 | 0.33 |
|  | Cell Mean | 0.00 | 0.00 | 0.00 | 0.33 | 0.67 | 0.67 | 1.00 | 1.00 | 1.00 | 1.00 | 1.00 | 0.67 | 0.00 | 0.00 | 0.00 | 7.33 |  |
| Midline Ablation | 1 | 0.00 | 0.00 | 0.00 | 1.00 | 1.00 | 0.00 | 1.00 | 1.00 | 1.00 | 1.00 | 0.00 | 0.00 | 0.00 | 0.00 | 0.00 | 6 | 0.40 |
|  | 2 | 0.00 | 0.00 | 0.00 | 0.00 | 0.00 | 0.00 | 0.00 | 1.00 | 1.00 | 1.00 | 1.00 | 0.00 | 0.00 | 0.00 | 0.00 | 4 | 0.27 |
|  | 3 | 0.00 | 0.00 | 0.00 | 0.00 | 0.00 | 0.00 | 0.00 | 0.00 | 0.00 | 0.00 | 0.00 | 0.00 | 0.00 | 0.00 | 0.00 | 0 | 0.00 |
|  | 4 | 0.00 | 0.00 | 0.00 | 0.00 | 0.00 | 0.00 | 0.00 | 0.00 | 0.00 | 0.00 | 0.00 | 0.00 | 0.00 | 0.00 | 0.00 | 0 | 0.00 |
|  | Cell Mean | 0.00 | 0.00 | 0.00 | 0.25 | 0.25 | 0.00 | 0.25 | 0.50 | 0.50 | 0.50 | 0.25 | 0.00 | 0.00 | 0.00 | 0.00 | 2.50 |  |

